# Altered polarization of PAR-2 signaling during airway epithelial remodeling

**DOI:** 10.1101/2020.01.09.900555

**Authors:** Ryan M. Carey, Jenna R. Freund, Benjamin M. Hariri, Nithin D. Adappa, James N. Palmer, Robert J. Lee

## Abstract

**Background:** Protease-activated receptor 2 (PAR-2) is activated by proteases involved in allergy and triggers airway epithelial secretion and inflammation. PAR-2 is normally expressed basolaterally in differentiated nasal ciliated cells.

**Objective:** We tested if epithelial remodeling during diseases characterized by loss of cilia and squamous metaplasia may alter PAR-2 polarization.

**Methods:** Endogenous PAR-2 responses were measured by live cell calcium and cilia imaging, measurement of fluid secretion, and quantification of cytokines. We utilized airway squamous cell lines, primary differentiated air-liquid interface cultures, and tissue explants. Cells were exposed to disease-related modifiers that alter epithelial morphology, including IL-13, cigarette smoke condensate, and retinoic acid deficiency. We used concentrations and exposure times that altered epithelial morphology without causing breakdown of the epithelial barrier, likely reflecting early disease states.

**Results:** PAR-2 signaling in airway squamous cells activated calcium and inflammatory responses. Squamous cells cultured at air liquid interface (ALI) responded to PAR-2 agonists applied both apically and basolaterally. Primary well-differentiated nasal epithelial ALI cultures responded only to basolateral PAR-2 stimulation. Primary cultures exposed to IL-13, cigarette smoke condensate, or reduced retinoic acid responded to both apical and basolateral PAR-2 stimulation. Nasal polyp tissue, but not control middle turbinate, exhibited apical calcium responses to PAR-2 stimulation. However, isolated ciliated cells from both polyp and turbinate maintained basolateral PAR-2 polarization.

**Conclusions:** Squamous metaplasia and/or loss of cilia enhances apical PAR-2 responses. Altered PAR-2 polarization in dedifferentiated or remodeled epithelia may contribute to increased sensitivity to inhaled protease allergens in inflammatory airway diseases.

## INTRODUCTION

Protease-activated receptors (PARs) are G protein-coupled receptors (GPCRs) that can drive inflammation in asthma, chronic rhinosinusitis (CRS), and allergic rhinitis.^1, 2^ PARs are activated by proteolytic cleavage of the extracellular N-terminus, exposing an intramolecular tethered ligand.^1, 2^ PAR-2 may be activated by mast cell tryptase,^3^ neutrophil elastase,^4^ *Alternaria* fungal proteases,^5^ and house dust mite proteases^6^ and may promote Th_2_ inflammation in bronchial epithelial cells.^7^ We previously found that PAR-2 is expressed on the basolateral membrane of healthy well-polarized differentiated primary sinonasal epithelial ciliated cells, where it elevates intracellular calcium to increase ciliary beating and apical membrane Cl^-^ secreation,^8^ suggesting PAR-2 promotes epithelial mucociliary clearance in response to protease activity.

We hypothesized basolateral polarization of PAR-2 may limit its activation except during times of epithelial barrier breakdown during infection or inflammation. Alteration of epithelial polarization or cell type composition occurs in obstructive airway diseases such as asthma and COPD.^9, 10^ Loss of airway cilia may occur with viral or bacterial infection,^11-14^ type-2 inflammation-driven remodeling,^15^ or smoking.^16, 17^ Squamous metaplasia occurs after bacterial infection, particularly in cystic fibrosis-related CRS,^18^ and is frequently reported in CRS patient tissue samples.^19, 20^

Much of the literature on airway remodeling focuses on barrier dysfunction, but airway cells *in vitro* can maintain tight junctions and barrier function despite substantial morphological changes, including loss of cilia.^12^ Less is known about how signaling is altered before large scale barrier breakdown in severe disease. We hypothesized that loss of epithelial polarization may affect PAR-2 signaling independently of barrier disruption, perhaps sensitizing cells to protease allergens.

We first tested activation of PAR-2 by *Aspergillus fumigatus*, the most prevalent fungal cause of pulmonary allergic disease, including allergic bronchopulmonary aspergillosis associated with chronic lung injury in patients with cystic fibrosis (CF) or chronic asthma.^21^ We next examined PAR-2 signaling in airway squamous cells using pharmacology and live cell imaging. We then tested PAR-2 signaling in well-differentiated air-liquid interface (ALI) cultures of primary nasal cells exposed to disease-related modifiers, including IL-13, cigarette smoke condensate (CSC), and retinoic acid deficiency, all of which alter epithelial morphology. We also examined isolated tissue from inflamed nasal polyp and compared it with control middle turbinate. The data below reveal important alterations of the polarity of PAR-2 signaling in squamous cells and primary epithelial cells de-differentiated through disease-relevant mechanisms.

## METHODS

### Materials

A detailed list of materials used are in the Supplementary Methods. Unless indicated, all other regents were from Sigma Aldrich (St. Louis, MO USA).

### General experimental methods

Cell line culture was as described.^8, 22, 23^ Fungal culture was carried out as described.^24^ Imaging of ASL height was performed as described.^25^ FITC-conjugated dextran permeability was measured as described.^26^ Glucose permeability was measured as described.^27^ Calcium imaging was performed as described.^8, 22, 23, 28^ CBF was measured using the Sisson-Ammons Video Analysis sytem^29^ as described.^8, 23^ Further details are provided in the **Supplementary Methods**.

### Generation of primary sinonasal ALI cultures (ALIs) from residual surgical material

Tissue acquisition was carried out in accordance with The University of Pennsylvania guidelines regarding use of residual clinical material, using tissue from patients ≥18 years of age undergoing surgery for sinonasal disease (CRS) or other procedures (e.g. trans-nasal approaches to the skull base). Institutional review board approval (#800614) and written informed consent was obtained in accordance with the U.S. Department of Health and Human Services code of federal regulation Title 45 CFR 46.116 and the Declaration of Helsinki. Clinical information regarding patient samples is listed in **Supplementary Table S1**. Experiments using primary mouse nasal septal ALI cultures utilized residual tissue from mice euthanized for other experimental purposes according to the “Guide for the Care and Use of Laboratory Animals” (Institute of Laboratory Animal Resources, National Research Council) with institutional approval. Cell culture was carried out as described.^8, 22, 23^ More details are in the **Supplementary Methods**.

### Data and statistical analysis

Images were acquired in Metamorph, Metafluor, and/or Micromanager^30^ and analyzed in FIJI.^31^ Multiple comparisons were done in GraphPad Prism (La Jolla, CA) using one-way ANOVA with Bonferroni (for pre-selected pairwise comparisons), Tukey-Kramer (for comparing all values), or Dunnett’s (for comparing to control value) posttests; *p* <0.05 was considered statistically significant. All data are mean ± SEM.

## RESULTS

### *Aspergillus fumigatus* conditioned media can directly activate PAR-2

While *Alternaria* proteases are well documented to activate PAR-2,^32, 33^ evidence exists that *Aspergillus fumigatus* also activates PAR-2 in the airway^7^ and cornea.^34^ *A. fumigatus* is a common cause of allergic bronchopulmonary aspergillosis or allergic fungal rhinosinusitis,^21^ and can cause fatal invasive aspergillosis.^35^ To confirm if *A. fumigatus* produces PAR-2-activating proteases, we utilized a fluorogenic assay (Trio assay) to directly visualize PAR-2 activation by its interaction with β-arrestin,^36^ which associates with activated and phosphorylated GPCRs and mediates further signal transduction and/or internalization to endosomes^37^ (**Fig 1A**). This assay is based on a tripartite GFP, with the 11^th^ β strand fused to the C-terminus of PAR-2, the 10^th^ β-strand fused to β-arrestin, and β strands 1-9 expressed as a soluble protein.^36^ When PAR-2 is activated, association with β-arrestin allows a complete GFP to form and increases fluorescence (**Fig 1B**). This approach eliminates other effects that may impinge on downstream signaling (e.g., toll-like receptor activation during exposure to fungal conditioned media).

**FIG 1.**
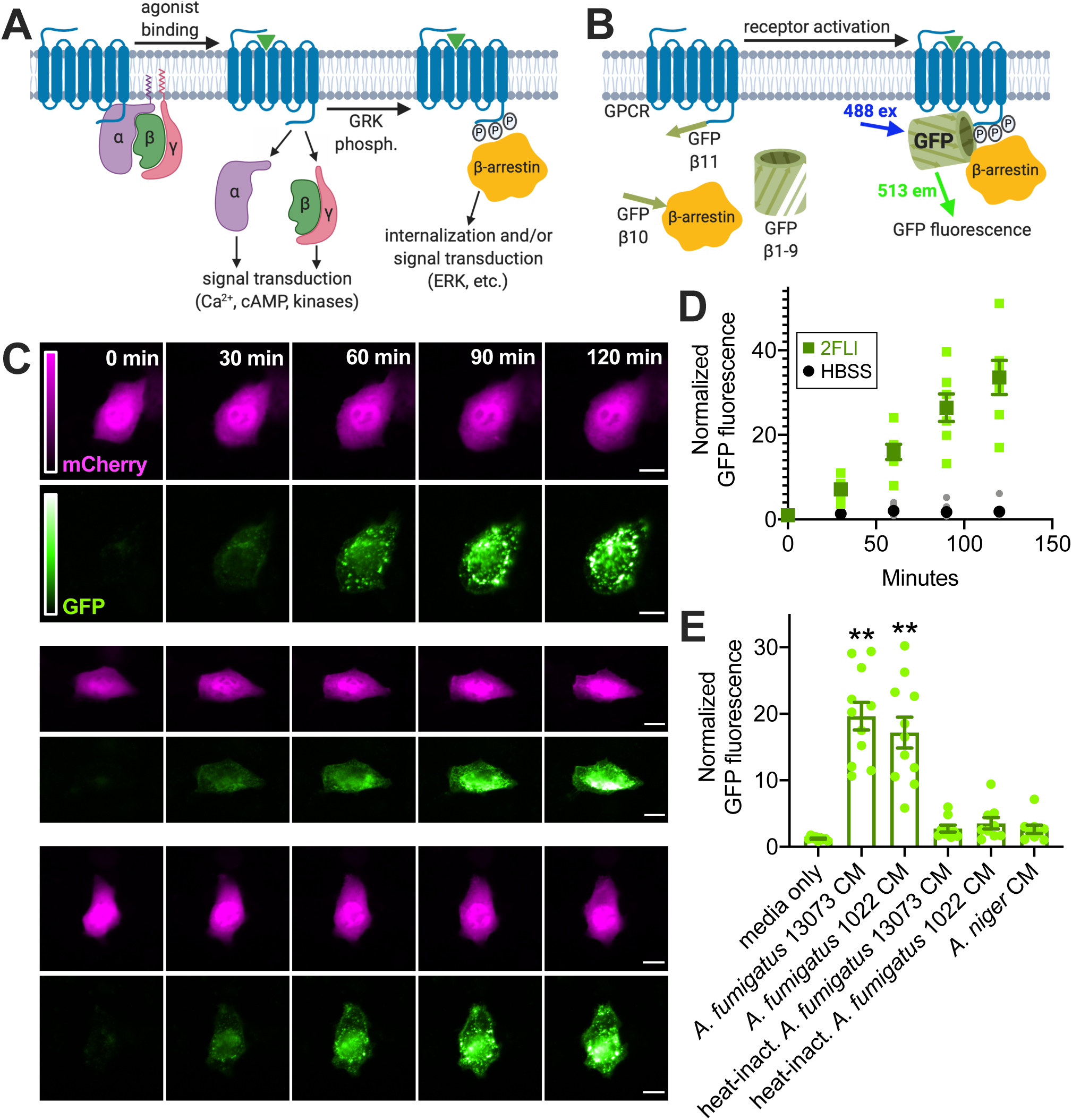
*Aspergillus fumigatus* conditioned-media (CM) directly activates PAR-2 in airway cells. **A**, Diagram of G protein-coupled receptor (GPCR) interaction with β-arrestin after activation and phosphorylation by G receptor kinase (GRK) **B**, Diagram of the Trio assay^36^ used to detect PAR-2 activation. Association of the PAR-2 with β-arrestin allows the formation of a complete fluorescent GFP molecule. **C**, Representative images of A549 cells expressing PAR-2 Trio components (plus mCherry as transfection control) stimulated with 10 µM 2FLI for 0-120 min. Scale bar is 15 µm. **D**, Normalized GFP fluorescence in cells stimulated with 2FLI or buffer alone (Hank’s balanced salt solution [HBSS]) at time points taken over 2 hours. **E**, Normalized GFP fluorescence in A549 cells stimulated for 10 min with 25% fungal CM or media (diluted in HBSS) as indicated then washed with HBSS and observed at 90 min after the fungal stimulation. Representative images are in the Figs E1 and E2. Significance in *E* determined by 1-way ANOVA with Dunnett’s posttest comparing values to media only control; \*\**p* <0.01. Data points in *D* and *E* are independent experiments imaged on different days (n = 5-9). Figures in *A* and *B* created with Biorender.com.

We visualized PAR-2 activation via appearance of GFP fluorescence A549 alveolar cells (**Fig 1 and Supplementary Fig S1**) and Beas-2B bronchial cells (**Supplementary Fig S2**). Stimulation with peptide PAR-2 agonist 2FLI resulted in appearance of punctate GFP fluorescence over the course of 2 hours (**Fig 1C-D and Supplementary Fig S2A-B**). In cells imaged at 90 min, 2FLI increased GFP fluorescence ∼30-fold while PAR-4 agonist AY-NH_2_ had no effect (**Supplementary Fig S1A and S2C**). Exposure to conditioned media (CM) from *A. fumigatus*, but not *A. niger*, increased GFP fluorescence (**Fig 1E and Supplementary Figs S1B and S2G-K**), signaling PAR-2 activation. Heat inactivation of the CM (100 °C for 20 min) eliminated PAR-2 activation (**Fig 1E and Supplementary Figs S1B, S2H and S2K**). No activation of the β-2 adrenergic receptor was observed (**Supplementary Fig S1C**). These results support that *A. fumigatus* activates of PAR-2, likely via a secreted heat labile protease.

Controversy exists over whether certain host and pathogen proteases activate PAR-2 by cleavage at the activating site or inhibit PAR-2 by cleavage downstream, removing the tethered ligand and “disarming” it.^38^ Neutrophil and *Pseudomonas aeruginosa* elastase have both been suggested to either activate^4, 39, 40^ or disarm PAR-2.^41, 42^ Differential results may be due to altered protein processing in different cell types or different assays used. We tested if several proteases implicated in allergy can activate PAR-2 expressed in airway cells, and observed activation with human lung tryptase, neutrophil elastase, and dust mite protease Der p 3 (**Supplementary Fig S2L-R**).

### PAR-2 is expressed and functional in squamous airway epithelial cells

We observed expression of PAR-2 in nasal septal squamous RPMI 2650 cells (**Supplementary Fig S3**). PAR-2 peptide agonist 2FLI increased intracellular calcium (**Supplementary Fig S4**) and transiently activated protein kinase C (**Supplementary Fig S5**) in both RPMI 2650 and lung squamous line NCI-H520. We also observed activation of PAR-2 by human lung tryptase and Der p 3 (**Supplementary Fig S6**). PAR-2 activation also had a small but significant effect of acutely reducing squamous cell line metabolism and/or proliferation through lowering of cAMP, but did not activate apoptosis (**Supplementary Fig S7**). PAR-2 activation increased secretion of TNFα and GM-CSF from RPMI 2650 cells and IL-6 and GM-CSF from NCI-H520 cells, likely involving dual coupling to G_q_ and G_i_ G protein isoforms (**Supplementary Fig S8**). Thus, PAR-2 in airway squamous cells can regulate inflammation. We next tested how PAR-2 signaling and polarization change with epithelial de-differentiation and/or remodeling using primary cells.

### Primary airway epithelial ciliated air liquid interface (ALI) cultures can be dedifferentiated/remodeled by exposure to a type II cytokine or submersion

To begin to understand how epithelial remodeling changes polarization of PAR-2 in primary sinonasal cells, we initially de-differentiated/remodeled primary ciliated air-liquid interface (ALI) cultures by two methods. The first was exposure to IL-13 (10 ng/ml, basolateral), which induces cilia loss and goblet cell metaplasia.^43, 44^ The second was apical submersion, which induces a more squamous phenotype.^43, 45^ ALIs were cultured for 21 days to establish a well-ciliated baseline (**Fig 2A**). IL-13 or apical submersion both resulted in a decrease in the number of ciliated cells over 6 days, shown by a reduction in β-tubulin IV immunofluorescence (**Fig 2A-B**). While IL-13 increased goblet cell specific Muc5AC intensity (**Figs 2A and C**), submersion only minimally increased Muc5AC at 6 days (**Figs 2A and C**). Submersion instead increased expression of transgutaminase 1 (TG-1; measured via ELISA), previously reported to reflect squamous differentiation^46, 47^ (**Fig 2D**). Transepithelial electrical resistance (TEER) remained intact up to 4 days (**Fig 2E**), and maintenance of epithelial barrier function was confirmed by measurement of FITC-conjugated dextran flux at day 4 as well as ability of cells to maintain low apical glucose levels (**Supplementary Fig S9A**). Evidence of cell death (LDH release) was not observed (**Supplementary Fig S9B**).

**FIG 2.**
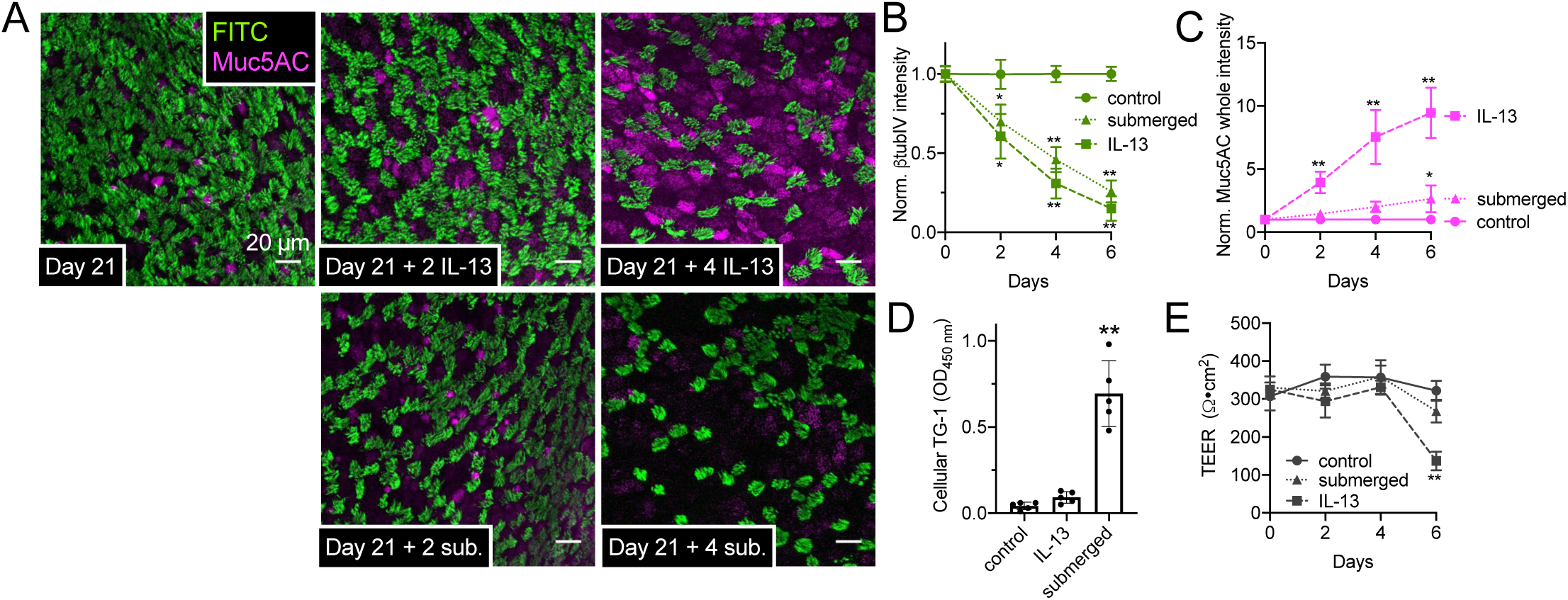
Loss of cilia with IL-13 treatment or submersion of air liquid interface (ALI) cultures of primary human nasal cells. **A**, Representative immunofluorescence (IF) of cilia (β-tubulin IV; green) and Muc5AC (magenta) in ALIs 21 days after air (top left). Top middle and right show cilia loss after further 2-4 days basolateral IL-13. Bottom middle and right show loss of cilia with less Muc5AC with apical submersion. **B and C**, Cilia loss (*B*; normalized β-tubulin IV IF) and Muc5AC increase (*C*) in cultures (starting at day 21) exposed to subsequent IL-13, apical submersion, or no treatment (control). Ten fields from one ALI from one patient were imaged and averaged for an independent experiment; results shown are mean ± SEM of 4-6 independent experiments (4-6 patients). **D**, Squamous marker TG-1 quantified by ELISA at day 25 (21 days at air then 4 subsequent days of IL-13, submersion, or no treatment). Each data point is an independent ALI from a different patient (n = 5 total ALIs). Significance by 1-way ANOVA with Dunnett posttest; \*\**p*<0.01. **E**, Transepithelial electrical resistance (TEER) at time points as in *B* and *C*. Significance by 1-way ANOVA with Bonferronni posttest; \**p*<0.05 and \*\**p*<0.01.

We thus chose the 4-day exposure time to IL-13 or submersion (21 days of differentiation plus further 4 days of IL-13 or submersion, here termed Day 21 + 4) for further experiments as it was a timepoint where the epithelium was remodeled but barrier function remained intact. At this time point, we saw no differences in expression of PAR-2 by qPCR (**Supplementary Fig S9C**), which mimicked a lack of differences observed in nasal polyp tissue (frequently characterized by type II inflammation) and control middle turbinate tissue (**Supplementary Fig S9C**), in contrast to another study.^48^ As a control, we did see increased expression of PAR-2 in ALIs after lipopolysaccharide (LPS), agreeing with previous results from bronchial fibroblasts (**Supplementary Fig S9C**).^49^

### Epithelial remodeling is associated with altered polarization of PAR-2 signaling

In primary ALIs at 5 days after seeding, when ALIs have electrical resistance but cilia have not yet developed, both apical or basolateral 2FLI or trypsin induced calcium responses (**Fig 3A and F**). After a saturating dose of apical 2FLI, another increase in apical 2FLI did not elicit a further response, while application of basolateral 2FLI did (**Fig 3A**, left). This suggests that 2 separate pools of PAR-2 receptors exist, separated by the apical-to-basolateral tight junction diffusional barrier. By day 21, responses to apical 2FLI or trypsin were lost, while responses to basolateral stimulation were intact (**Fig 3B and F**).

**FIG 3.**
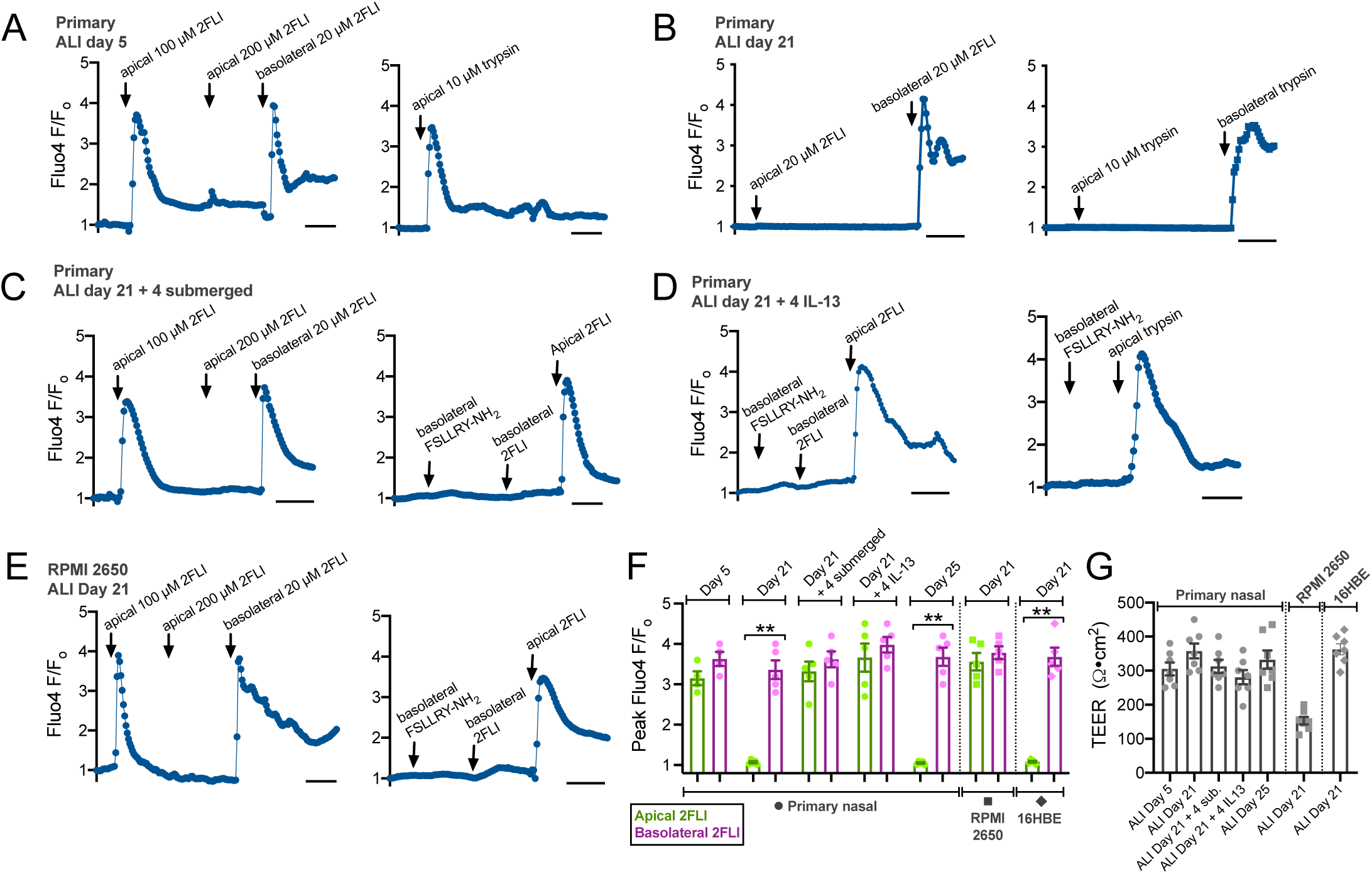
Polarization of PAR-2 signaling in ALIs. **A-D**, Primary human nasal ALIs loaded with Fluo-4 were stimulated apically or basolaterally with PAR-2 agonist 2FLI or trypsin. ALIs at day 5 (before ciliogenesis) responded to apical 2FLI (*A*, left) or trypsin (*A*, right). Basolateral 2FLI elicited further response after apical saturation, suggesting two physically separate PAR-2 pools (*A*, left). Responses to apical 2FLI or trypsin were lost by day 21 (*B*) but regained if after subsequent 4 days of submersion (*C*) or basolateral IL-13 (*D*). Apical responses to 2FLI were observed even with basolateral PAR-2 antagonist (FSLLRY-NH_2_; 100 µM), supporting distinct PAR-2 pools. Representative traces shown from single ALIs. **E**, RPMI2650 squamous ALIs exhibited responses to apical 2FLI. Subsequent application of basolateral 2FLI or inhibition of basolateral PAR-2 with FSLLRY-NH_2_ also suggest physically separated apical vs basolateral PAR-2 pools. **F**, Quantification of peak Fluo-4 F/F_o_ from experiments as in *A-E* (mean ± SEM; 5-7 experiments using ALIs each different patients). Significance by 1-way ANOVA with Bonferroni posttest comparing apical vs basolateral 2FLI; \*\**p*<0.01. **G**, TEER from cultures used in *A-F* confirmed 4 days of submersion or IL13 did not disrupt the epithelial barrier. Significance by 1-way ANOVA with Bonferroni posttest.

If primary cells were incubated for a further 4 days with the apical side submerged (Day 21 + 4 submerged) to promote squamous de-differentiation, the response to apical 2FLI was again observed (**Fig 3C and F**). Basolateral application of antagonist FSLLRY-NH_2_ blocked the response to basolateral but not apical 2FLI (**Fig 3C**, right), ruling out 2FLI diffusion across the epithelial barrier to the basolateral side. Similar results were seen at ALIs after 4 subsequent days of basolateral IL-13 (**Fig 3D**). If primary ALIs were carried out to day 25 with no IL-13 and no apical submersion, only basolateral 2FLI responses were observed (**Fig 3F**). RPMI2650 squamous cells cultured at ALI also had 2FLI responses with application to either the basolateral or apical side; basolateral addition of antagonist blocked basolateral but not apical responses (**Fig 3E and F**). No significant differences in TEERs were observed (**Fig 3G**).

### Altered PAR-2 polarization increases epithelial fluid secretion during apical exposure to a PAR-2 agonist

We measured airway surface liquid (ASL) height in primary ALIs. Physiological ASL was labeled with Texas red dextran sonicated in perfluorocarbon (PFC) and ASL height was measured from orthogonal confocal slices. Primary ALIs were compared at differentiation day 25 (**Fig 4A**) or after 21 days + 4 subsequent days IL-13 (**Fig 4B**). Baseline ASL height was not different. Basolateral 2FLI caused an increase in ASL height in both groups (**Figs 4A-C**) that was inhibited by basolateral application of the Na^+^K^+^2Cl^-^ co-transporter (NKCC) inhibitor bumetanide (**Fig 4C**), showing that this reflected fluid secretion. However, when 2FLI was added to the apical side in PFC, IL-13 treated cultures, but not control cultures, exhibited increased ASL height (**Figs 4A-C**) that was blocked by calcium-activated Cl^-^ channel (CaCC) inhibitor CaCC_inh_-A01 but not CFTR_inh_-172 (**Figs 4B-C**).

**FIG 4.**
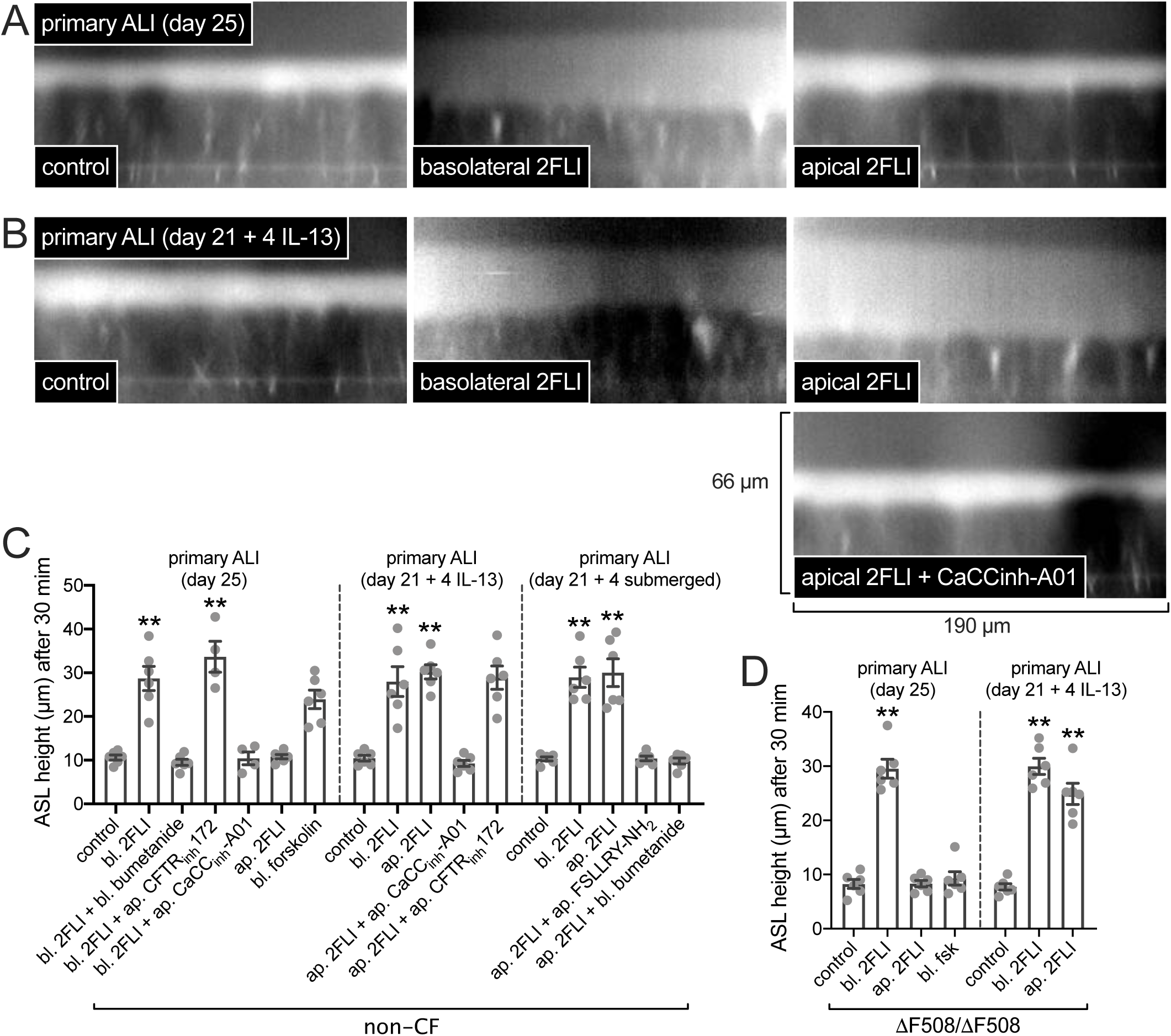
Epithelial remodeling is associated with enhanced fluid secretion to apical PAR-2 stimulation. **A**, Representative orthogonal confocal sections of control cultures with Texas red dextran-labeled airway surface liquid (ASL) showing increase in ASL height (reflecting fluid secretion) with basolateral but not apical 2FLI (∼25 µM). **B**, Representative sections of IL-13-treated cultures showing increased secretion with either basolateral or apical 2FLI and inhibition by calcium-activated chloride channel (CaCC) inhibitor CaCC_inh_-A01 (∼10 µM). **C**, Peak ASL heights from independent experiments, each the average of 10 ASL measurements from one ALI; 6 experiments per condition. Note no inhibition of secretion with CFTR_inh_172 (∼10 µM), but inhibition with NKCC1 inhibitor bumetanide (100 µM) or PAR-2 antagonist FSLLRY-NH_2_ (∼10 µM). Drugs applied to the apical side were sonicated in perfluorocarbon to not disturb the aqueous ASL layer, thus apical concentrations are approximate. **D**, Data experiments using ΔF508/ΔF508 homozygous CF ALIs. Note no response in CF cultures to basolateral forskolin (20 µM), which elevates cAMP and activates CFTR (non-CF cultures responded in *C*). Each point is an independent experiment using cells from one patient (5 patients total). Significance in *C* and *D* by 1-way ANOVA with Dunnett’s posttest.

Similar responses to apical 2FLI were observed after 21 days of differentiation + 4 days of submersion (**Fig 4C**). Apical 2FLI response was blocked by PAR-2 antagonist FSLLRY-NH_2_ (**Fig 4C**). We performed experiments using cells from cystic fibrosis (CF) patients homozygous for ΔF508 CFTR. CF ALIs robustly secreted fluid in response to PAR-2 stimulation. These data suggest that apical PAR-2 agonist exposure does not result in epithelial fluid secretion in well differentiated cultures, but remodeling or dedifferentiation results in apical PAR-2-activated fluid secretion via CaCC.

### Epithelial remodeling and altered PAR-2 polarization enhances cytokine secretion during apical exposure to PAR-2 agonists

To further examine effects of PAR-2 signaling alterations above, we examined secretion of TGF-β2, an important epithelial-derived cytokine in type 2 airway diseases,^50^ In RPMI 2650 ALIs, TGF-β2 secretion was activated by basolateral 2FLI and inhibited ∼50% by G_i_ inhibitor pertussis toxin (PTX; **Fig 5A**). In contrast, activation of the TNF-α receptor, which is not a GPCR, increased TGF-β2 that was not inhibited by PTX (**Fig 5A**). In primary differentiated ALIs, apical 2FLI had no effect, but basolateral 2FLI increased TGF-β2 that was inhibited by basolateral but not apical FSLLRY-NH_2_ (**Fig 5B**). Similarly, apical trypsin or lung tryptase had little effect while basolateral trypsin and tryptase increased TGF-β2 secretion (**Fig 5B**).

**FIG 5.**
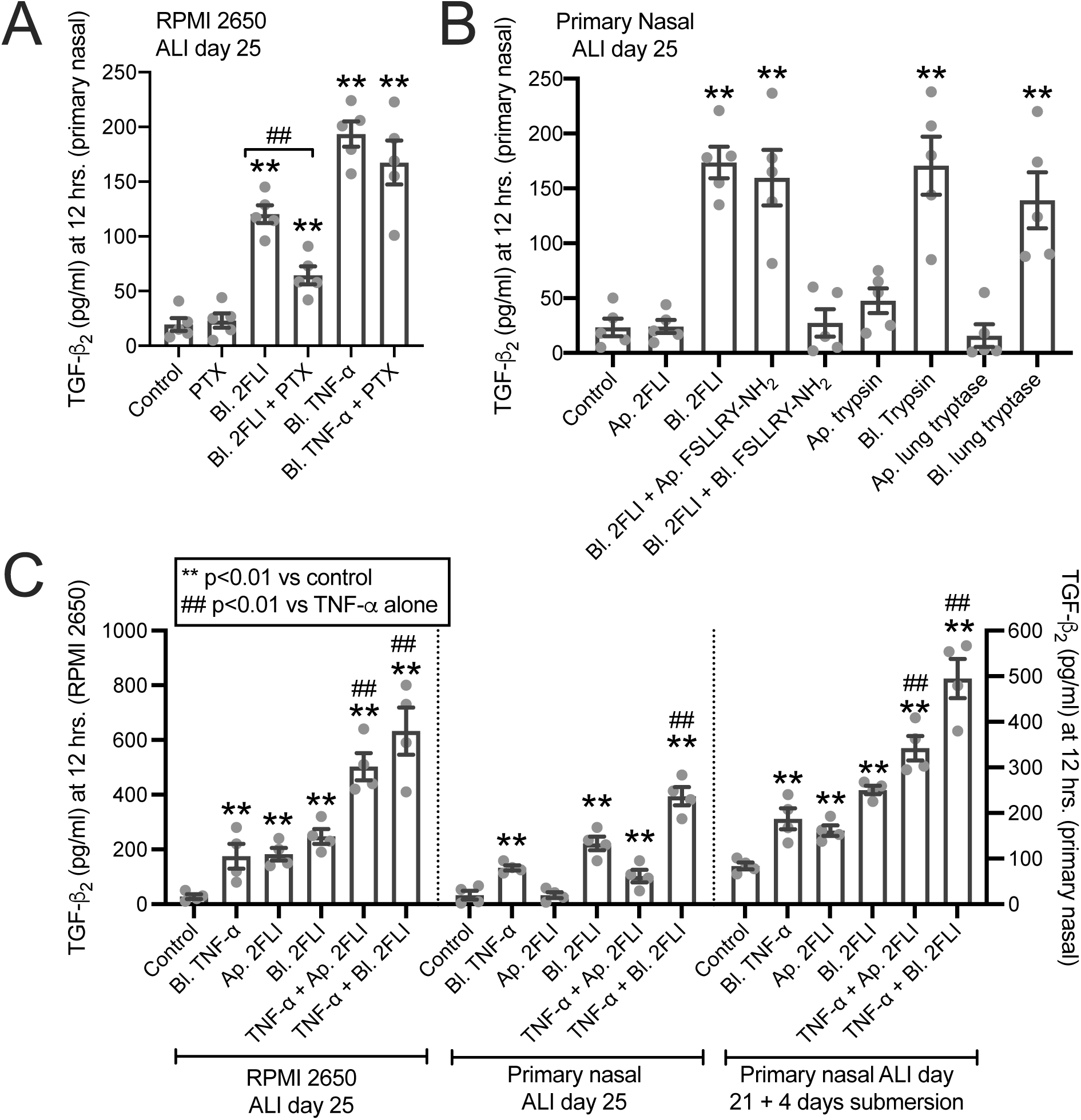
Altered PAR-2 polarity translates to altered TGF-β2 secretion with PAR-2 stimulation both alone and in combination with TNFα. **A**, Bar graph of RPMI 2650 ALI TGF-β2 secretion with basolateral (bl.) application of 2FLI (20 µM) or TNFα (10 ng/ml) ± pertussis toxin (100 ng/ml; 18 hrs pretreatment). **B**, Bar graph of primary nasal ALIs showing TGF-β2 secretion with basolateral (bl.) but not apical (ap.) 2FLI (20 µM), trypsin (10 nM), or tryptase (10 nM). **C**, Bar graph showing TGF-β2 secretion with apical vs basolateral 2FLI and combinatorial effects with TNFα. RPMI 2650 squamous ALIs and primary ALIs exposed to 4 days of submersion to induce squamous differentiation exhibited apical 2FLI responses while control ALIs did not. Significance in *A-C* by one-way ANOVA with Dunnett’s posttest comparing values to respective control; **p <0.01. In *A*, ^##^p<0.01 between basolateral 2FLI and 2FLI with PTX by one-way ANOVA with Bonferroni posttest. In C, ^##^p<0.01 vs TNFα alone by one way ANOVA with Bonferroni posttest.

In RPMI 2650 ALIs or ALIs exposed to 4 days of submersion, apical 2FLI increased TGF-β2 (**Fig 5C**). It also potentiated TGF-β2 secretion in the presence of TNF-α (**Fig 5C**). In RPMI 2650 ALIs and primary ALIs treated with basolateral TNF-α, basolateral 2FLI caused a further increase of TGF-β2 beyond TNF-α alone (**Fig 5C**). In RPMI 2650s and ALIs exposed to 4 days of submersion, this occurred with apical 2FLI as well (**Fig 5C**). Similar results were observed in mouse nasal septum ALI cultures with responses lost in ALIs from PAR-2 knockout (par2^-/-^) mice (**Supplementary Fig S10**).

### Impaired ciliogenesis resulting from cigarette smoke condensate (CSC) exposure or retinoic acid deficiency also correlates with altered polarization of PAR-2

The above data suggest that epithelial de-differentiation or remodeling affects the polarization of PAR-2 and may sensitize epithelial cells to protease allergens by allowing apical PAR-2 activation. We tested if other alterations of epithelial development and ciliogenesis cause this phenotype. We used cigarette smoke condensate (CSC), shown to inhibit ciliogenesis in nasal ALIs.^17^ Human ALIs cultured in the presence of 10-30 µg/ml CSC exhibited impaired ciliogenesis at day 20 (**Supplementary Fig S11A-B**). Higher concentrations increased epithelial permeability, but barrier function remained intact up to 30 µg/ml CSC (**Supplementary Fig S11C-D**).

In day 20 human nasal ALIs cultured in 10-30 µg/ml CSC, apical 2FLI, Der 3 p, and tryptase induced calcium responses, while control cells (0 µg/ml CSC) did not respond (**Supplementary Fig S11E**). Apical PAR-2 activation also induced TGF-β2 and GM-CSF in human nasal ALIs cultured in 10-30 µg/ml CSC, while control cells responded only to basolateral PAR-2 stimluation (**Supplementary Fig S11E-F**). Similar results were observed in mouse ALIs, and par-2^-/-^ mice confirmed responses were indeed due to PAR-2 (**Supplementary Fig S12**).

The active metabolite of vitamin A, retinoic acid (R.A.), is involved in transcriptional control of ciliogenesis.^46^ R.A. has anti-allergic effects in mice^51, 52^ and *in vitro* cell models. Growing primary airway epithelial cells in the absence of R.A. induces a squamous phenotype.^53^ Cilia differentiation was measured by acetylated tubulin ELISA in ALIs grown in 50 nM (normal level), 15 nM, or no R.A. for 20 days. Squamous differentiation was measured by TG-1 ELISA. Retinoic acid reduced cellular TG-1 (**Supplementary Fig S13A**), increased acetylated tubulin (**Supplementary Fig S13B**), and increased TEER (**Supplementary Fig S13C**). R.A.-deficient (15 nM) ALIs exhibited apical 2FLI calcium responses (**Supplementary Fig S13D**) while R.A. replete (50 nM) ALIs did not (**Supplementary Fig S13D**). Similar results were observed with tryptase and PAR-2 agonist AC-5541 (**Supplementary Fig S13E**). Calcium responses correlated with GM-CSF secretion at 24 hrs. Thus, impairment of ciliogenesis by CSC or reduced R.A. can both alter epithelial polarity of PAR-2 signaling.

### Patient tissue explants from polyps, but not control turbinate, exhibit apical PAR-2 responses

To further test if epithelial remodeling observed in airway diseases translates to altered PAR-2 responses, we took tissue explants from nasal polyps, removed from CRS patients during functional endoscopic sinus surgery, and control middle turbinate tissue, removed from patients during the course of surgery for other indications (e.g., trans-nasal approaches to the skull base to remove pituitary tumors). Tissue was mounted, imaged by confocal microscopy, and stimulated apically with PAR-2 agonists. Polyp tissue exhibited apical responses to 2FLI, trypsin, or Der p 3, but not thrombin (**Fig 6A-B**). Control uninflamed middle turbinate tissue did not (**Fig 6C**). Data are summarized in (**Fig 6D**). Immunofluorescence revealed that PAR-2 was nonetheless basolaterally expressed in ciliated cells from nasal polyp (**Fig 6E**), as we previously reported for non-polyp tissue.^8^ Moreover, the calcium responses to PAR-2 stimulation were no enhanced in isolated polyp vs turbinate ciliated cells (**Fig 6F**). We thus hypothesized that apical PAR-2 responses do not originate from within the ciliated cells themselves.

**FIG 6.**
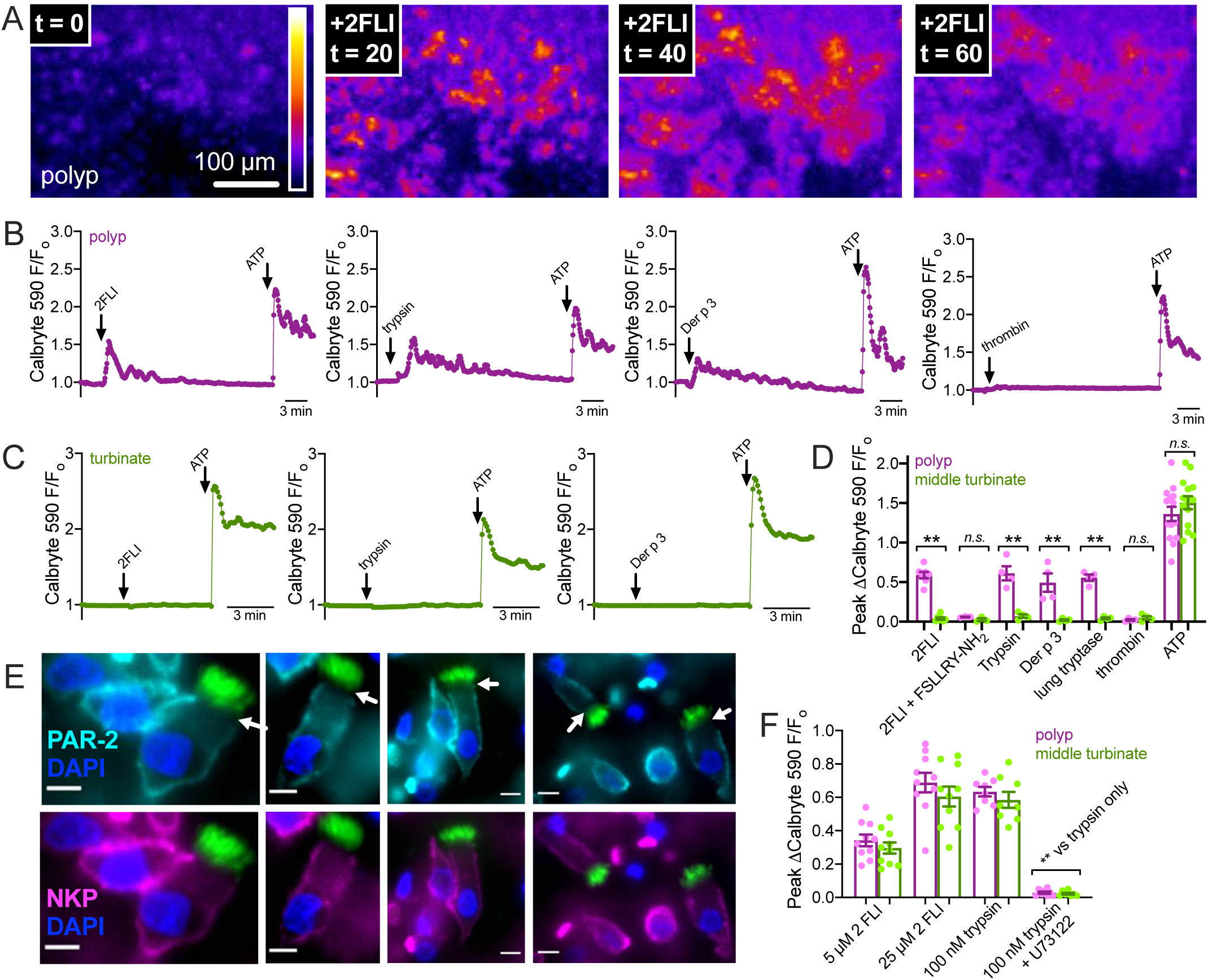
Nasal polyp exhibited calcium responses to apical PAR-2 stimulation while control turbinate did not. **A**, Representative image of 2-FLI induced calcium response (Calbryte 590) in a thin piece of polyp mucosa mounted in an Ussing chamber holder and imaged with a spinning disk confocal microscope. **B-C**, Representative traces of polyp (*B*) and turbinate (*C*) responses to apical 2FLI (50 µM), trypsin (1 µM), Der p 3 (1 µM), or thrombin (1 µM). ATP (100 µM), which activates apical purinergic receptors, was a positive control. **D**, Peak change in calcium in tissue treated apically with PAR-2 agonists. Each data point is an independent experiment using tissue from separate patients (n = 7 per condition). **E**, Immunofluorescence showing lack of apical PAR-2 staining in polyp ciliated cells with Na^+^K^+^ ATPase (NKP) as marker for the basolateral membrane and β-tubulin IV (cilia marker) shown in green. Pearson’s correlation and Mander’s overlap coefficients for NKP and PAR-2 in ciliated cells were both ≥0.95 in 10 images analyzed. **F**, Peak change in calcium in isolated ciliated cells; no significant difference observed between polyp and turbinate. Significance in *D* and *F* determined by one-way ANOVA with Bonferroni posttest; \*\**p*<0.01.

### Apical PAR-2 stimulation requires intracellular communication to increase ciliary beat frequency

To test if ciliated cells autonomously respond to apical PAR-2 stimulation, we measured ciliary beat frequency (CBF) to apical PAR-2 agonists in control and IL-13-treated cultures. We combined CBF measurement with two structurally distinct gap junction inhibitors, steroid-like small molecule inhibitor carbenoxolone (CBX) and peptide inhibitor Gap 27.^28^ Control cultures exhibited CBF increases with basolateral but not apical 2FLI; basolateral responses were unaffected by CBX (**Fig 7A, B, and G**). IL-13 treated cultures exhibited CBF increase with apical 2FLI (**Fig 7C**) inhibited by CBX or Gap 27 (**Fig 7D-F and G**). Basolateral 2FLI responses were not inhibited by CBX. Apical ATP responses were inhibited by neither CBX nor Gap 27 (**Fig 7G**). These data support a model by which well differentiated epithelium (**Fig 7H**) expresses basolateral PAR-2 within ciliated cells, as CBF increases do not require intercellular gap junction communication (**Fig 7H**). However, dedifferentiation of the epithelium results in apical expression of PAR-2 within non-ciliated cells that can increase CBF in ciliated cells only in the presence of functional gap junctions, likely to allow diffusion of calcium and/or IP_3_ (**Fig 7I**).

**FIG 7.**
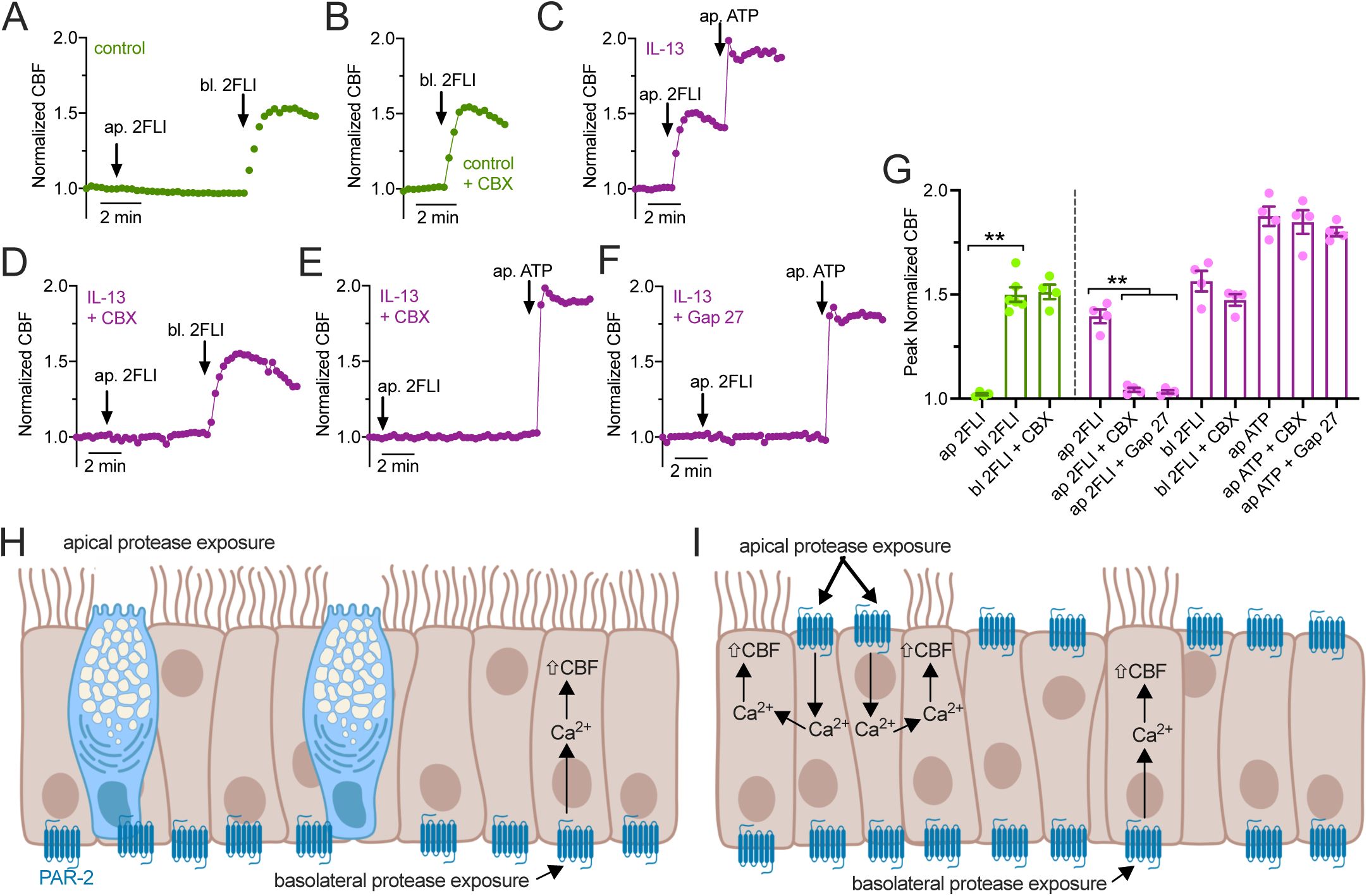
Ciliary beat frequency (CBF) measurements of nasal epithelial cells exposed to apical or basolateral 2FLI. **A-B**, Representative traces of CBF with 2FLI ± carbenoxolone (100 µM; 30 min preincubation). Control cultures (day 25 at ALI) exhibited no increased CBF with apical 2FLI application but a ∼50% increase in CBF with basolateral 2FLI. CBX did not inhibit effects of basolateral 2FLI. **C**, Representative trace of ∼50% increase in CBF with apical 2FLI in IL-13-treated ALI (21 days differentiation then 4 subsequent days with IL-13. **D-F**, Representative traces showing CBF in the presence of gap junction inhibitors CBX (*D* and *E*) or Gap 27 (*F*; 10 µM; 30 min pretreatment) in cultures treated with IL-13. Apical 2FLI responses were blocked while basolateral 2 FLI (*D*) or apical ATP (*E* and *F*) responses were intact. **G)** Bar graph showing Peak normalized CBF with agonist as used in A-F. Control cultures shown in green and IL-13 cultures shown in magenta. Each data point is an independent ALI culture from a separate individual patient (n = 5 per condition). Significance determined by 1-way ANOVA with Bonferroni posttest; **p<0.01 between bracketed values. **H-I**, Schematic of PAR-2 regulation of ciliary function in well-differentiated epithelium (modeled by control cultures; *H*) vs remodeled epithelium (modeled by IL-13 stimulated cultures; *I*). Data support that basolateral PAR-2 is expressed under both conditions and can regulate CBF directly within ciliated cells. Apical PAR-2 is not expressed in ciliated cells but apical PAR-2 stimulation can transmit signals (likely calcium, Ca^2+^) to ciliated cells through gap junctions. *H* and *I* created with Biorender.com.

## DISCUSSION

Prior literature on PAR-2 in the airway has focused on increased PAR-2 expression in inflamed airway mucosa.^2, 48, 54^ However, we show that an important overlooked consideration is the polarity of PAR-2, which can change in the absence of PAR-2 expression level changes. This may represent a novel mechanism by which the airway epithelium can be “sensitized” to apical exposure to inspired/inhaled proteases such as dust mite or *Alternaria* proteases. This may contribute to the pathogenesis of inflammatory airway diseases. We also confirm previous observations that PAR-2 is a player in detection of *A. fumigatus*.

Our data suggest that such alterations of epithelial morphology and composition due to inflammation, exposure to toxic compounds, or other environmental factors can all alter PAR-2 polarity (**Fig 8**). This may occur through increased Th2 cytokines, cigarette smoke exposure, or vitamin A deficiency. Exposure to inhaled allergens like dust mite or fungal proteases may normally be tolerated in healthy airway tissue if these allergens are cleared by mucociliary clearance before they can degrade the epithelial barrier and activate basolateral PAR-2. Moreover, airway submucosal gland secretions contain a variety of protease inhibitors that could neutralize pathogen proteases upon contact with mucus.^55, 56^ However, in some obstructive airway diseases, altered polarization of PAR-2 may combine with impaired protease inhibitor secretion due to gland duct plugging often observed.^57-59^ These phenotypes may result in epithelial sensitization to proteases at the apical membrane, increasing inflammatory responses.

**FIG 8.**
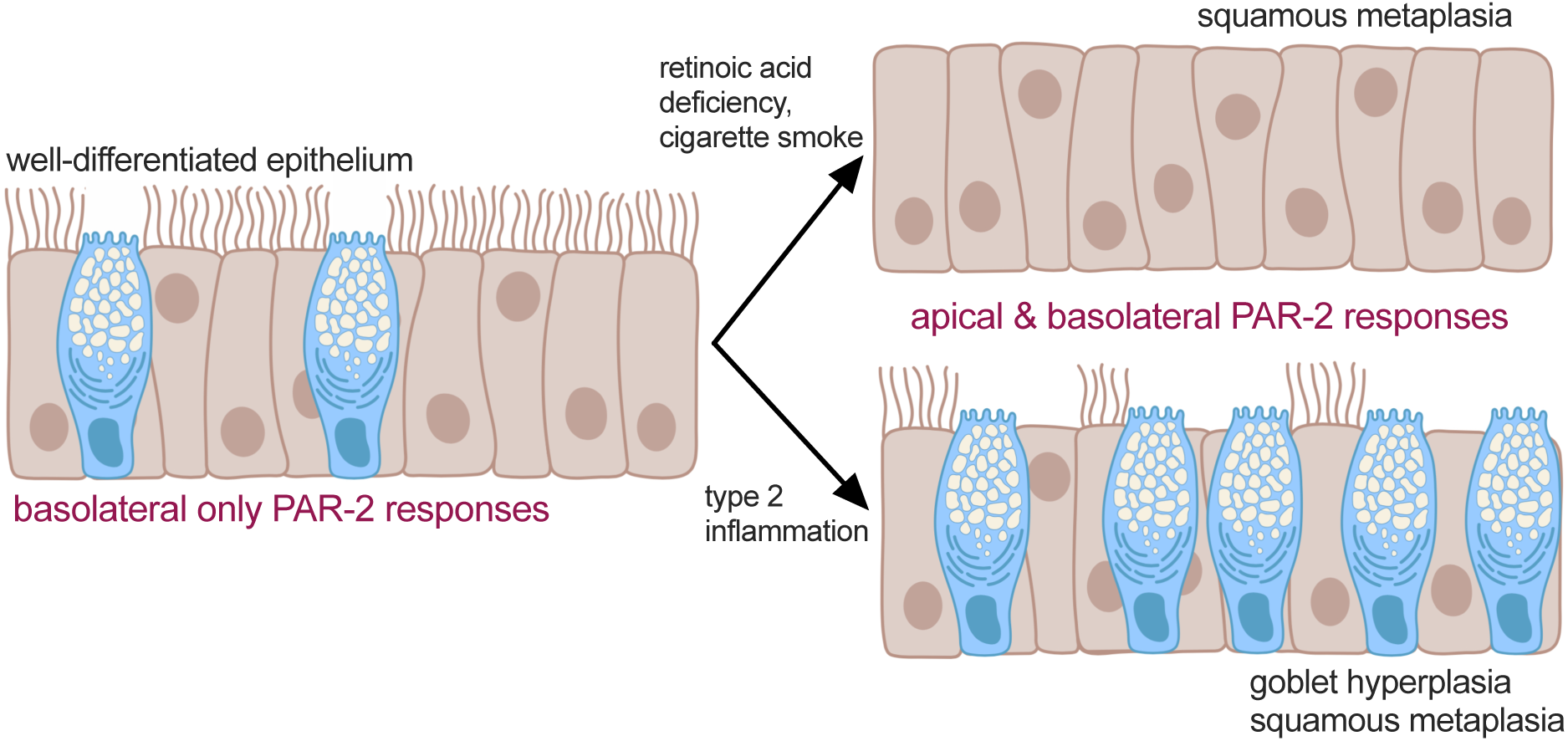
Model of altered PAR-2 signaling in airway disease. We hypothesize that well differentiated and highly ciliated airway epithelium can tolerate brief exposure to fungal or dust mite proteases due to the mainly basolateral expression of PAR-2, which is able to activate inflammatory and other responses only if the epithelium is breached. However, airway remodeling through a variety of disease modifiers can change the polarity of epithelial PAR-2 responses and may sensitize the epithelium to proteases and thus enhance inflammatory responses. Created with Biorender.com.

Squamous de-differentiation is a commonly observed histological finding in CRS patients,^19, 20^ CF patients,^60^ or smokers.^16, 61^ Squamous metaplasia also occurs during epithelial injury and repair.^15^ Prior studies have focused on the loss of ciliated cells and resulting reduced mucociliary clearance, but we show here that introduction of squamous cells into the epithelium can alter airway epithelial signaling, including an increase in apical PAR-2 responsiveness. This suggests disruption of the normal epithelial composition during disease has physiological consequences beyond reduced mucociliary clearance.

RPMI 2650 cells were proposed as a model of nasal permeability for drug transport studies^62, 63^ and cytokine release in allergy^64^ or viral infection.^65^ Our data here suggest that RPMI 2650 PAR-2 polarization is similar to squamous epithelial cells but not the same as well-differentiated ciliated cells, so interpretation of RPMI 2650 studies should be done cautiously. Altered polarization may extend to multi-drug resistance proteins or other transporters or channels relevant for epithelial absorption. RPMI 2650 studies might not reflect healthy sinonasal epithelium but may reflect inflamed remodeled epithelium such as polyp tissue.

Topical long-lasting PAR-2 antagonists might be useful in CRS patients with fungal or dust-mite activated allergic rhinitis by limiting apical PAR-2 activation, particularly if antagonists could be designed to not cross the epithelial barrier and block basolateral PAR-2 (e.g., a dextran conjugate) and delivered via topical nasal rinse. Such antagonists delivered via inhaler may be similarly useful in patients with allergic asthma. Alternatively, restoring a normal differentiated epithelial phenotype by reducing inflammation may lower the epithelial sensitivity to proteases.

## NONSTANDARD ABBREVIATIONS

ALI: air liquid interface
ASL: airway surface liquid
CaCC: calcium-activated chloride channel
CBF: ciliary beat frequency
CBX: carbenoxolone
CF: cystic fibrosis
CFTR: cystic fibrosis transmembrane conductance regulator
CM: conditioned media
COPD: chronic obstructive pulmonary disease
CRS: chronic rhinosinusitis
CSC: cigarette smoke condensate
ELISA: enzyme linked immunosubstrate assay
2FLI: 2-Furoyl-LIGRLO-NH_2_ PAR-2 agonist
GFP: green fluorescent protein
GM-CSF: granulocyte macrophage colony stimulating factor
GPCR: G protein-coupled receptor
GRK: G protein-coupled receptor kinase
IF: immunofluorescence
IL-13: interleukin 13
LDH: lactate dehydrogenase
MEM: minimal essential media
NKCC1: Na^+^K^+^2Cl^-^ cotransporter
PAR-2: protease-activated receptor 2
PFC: perfluorocarbon
TEER: transepithelial electrical resistance
TG-1: transglutaminase 1
TGF-β2: transforming growth factor β 2

## ACKNOWLEDGEMENTS

We thank Maureen Victoria (University of Pennsylvania) for technical assistance. We acknowledge Bei Chen (University of Pennsylvania) for isolating and propagating primary nasal cells used here. We thank Noam Cohen (University of Pennsylvania, Philadelphia VA Medical Center) for access to mouse nasal septum and cigarette smoke condensate and Laura Chandler (Philadelphia VA Medical Center) for fungal cultures. This study was supported in part by grants from the National Institutes of Health (R01DC016309 and R21AI137484) to R.J.L. The funders had no role in study design, data collection, analysis and interpretation of data, writing, or the decision to submit the article for publication. The article is solely the work of the authors and does not necessarily reflect the official views of the National Institutes of Health. There are no other conflicts of interest related to this research.

