## Supplementary Material for "Altered polarization of PAR-2 signaling during airway epithelial remodeling"

### Supplementary Methods

#### Materials

Fluo-4, CellRox, Texas Red dextran, FITC dextran, and AlexaFluor-labeled secondary donkey anti-mouse and donkey anti-rabbit were from ThermoFisher Scientfic (Waltham, MA USA). PAR-2 antibodies SAM-11 (mouse monoclonal) and EPR180953 (rabbit monoclonal), $Na^+/K^+$ ATPase (EP1845Y; rabbit monoclonal), and Glut1 (SPM498; mouse monoclonal) were from Abcam (Cambridge, UK). PAR-2 agonist peptide SLIGKV-$NH_2$ (Cat # H-4624) and scrambled LSIGKV-$NH_2$ (Cat # H-6428) were from Bachem (Torrance, CA USA). LDH assay kit (Cat # ab65393), 2-Furoyl-LIGRLO-$NH_2$ (2FLI; Cat #ab120800) was from Abcam. Glucose assay kit (#10009582), XTT assay kit (Cat #10010200), MTT assay kit (Cat #10009365) were from Cayman Chemical (Ann Arbor, MI USA). Multiple PAR-2 activating agonists were used in this study to ensure results were not an artifact of any one peptide or protease. Par-2 antagonist FSLLRY-$NH_2$ (Cat # 4751/1), Par-2 agonist SLIGRL-$NH_2$ (Cat # 1468/1; $EC_{50}$ 5 µM), scrambled LRGILS-$NH_2$ (Cat #3394/1), Par-4 agonist AY-$NH_2$ (Cat #1487/1), and Par-4 antagonist tcY-$NH_2$ (Cat # 1488) were from Tocris (Bristol, UK). AC-55541, a small molecule PAR-2 agonist ($EC_{50}$ = 6-7 µM), was from Cayman Chemical (Cat # 17736). Recombinant Der p 3 (Cat # MBS1096978) was from MyBioSource (San Diego, CA). ELISAs for GM-CSF (900-K30) and IL-6 (900-K16) were from Peprotech (Rocky Hill, NJ USA). Human TGFβ2 ELISA (DB250) and mouse TGFβ2 DuoSet ELISA (DY7346-05) were from R&D systems (Minneapolis, MN USA). Transglutaminase 1 (TG-1) ELISA (OKCD01601) was from Aviva Systems Bio (San Diego, CA USA). Acetylated tubulin ELISA (Cat # 7204) was from Cell Signaling Technologies (Danvers, MA USA). Cigarette smoke condensate was a gift of Noam Cohen (University of Pennsylvania) originally purchased from Murty Pharmaceuticals, Inc. (Lexington, KY; Batch R041018) as stock 40 mg/ml in DMSO stored at -80°C. Unless indicated below, all other regents were from Sigma Aldrich (St. Louis, MO USA).

#### Cell line culture

RPMI 2650 and NCI-H520 squamous cells as well as BEAS-2B and A549 cells were obtained from ATCC (Manassas, VA USA). 16HBE cells (SV-40 immortalized normal human bronchial) were obtained from D. Gruenert (University of California San Francisco, San Francisco, CA USA). Cells were grown in submersion in Minimal Essential Media with Earl's salts (Gibco; Gaithersburg, MD USA) supplemented with 10% FetalPlex (GeminiBioproducts; West Sacramento, CA USA) and 1%

penicillin/streptomycin mix (Gibco). RPMI 2650 and 16HBE cells were seeded for air liquid interface (ALI) culture onto Corning (Corning, NY) Transwells (0.3 $cm^2$ surface area; 0.4 um pore size) and grown for 5 days in the above media to confluence. Upon apical exposure to air, basolateral media was switched to primary cell differentiation media indicated below. ALIs were fed three times weekly. Transfection of submerged BEAS-2B and A549 cells was carried out in 8-well chamber slides (CellVis) with Lipofectamine 3000 as described.[1] Vectors for PAR-2 and β2 adrenergic receptor Trio assays were from Xiaokun Shu via Addgene. Cells were co-transfected with 2 pcDNA3 plasmids encoding the GFP β strands 1-9 (Addgene # 121684) and a pcDNA3 plasmid expressing the receptor fused to GFP β strand 11, an arrestin fused to GFP β strand 10, and mCherry as a transfection marker (Addgene #s 113609 or 113610) and imaged 24 hours after transfection at room temperature using standard GFP and TRITC filters on an Olympus IX83 microscope with 40x (0.75NA PlanFluor) objective. Sapphire CKAR or Sapphire CKAR(T/A) were from Jin Zhang via Addgene (Cat #s 118468 and 118469) and was imaged using the 380 channel of a fura-2 filter set (Chroma Technologies, Rockingham VT).

XTT assays were carried out as described in the manufacturer's instructions using phenol-free DMEM media (5.5 mM glc; GIBCO). Changes in XTT fluorescence were measured in real time in a Tecan Spark 10M plate reader equipped with heating (37°C) and gassing (5% $CO_2$). Technical replicates were used to ensure the reliability of single values. Experimental replicates presented in the text are independent experiments performed on different days with different batches of cells. PAR-2 immunofluorescence, biochemistry, and molecular biology in squamous cell lines were carried out as described for primary cells.[2]

**Fungal culture**

Cultures of *A. niger* (strain WB326 [ATCC16888] and *A. fumigatus* (NIH5233 [ATCC 13073] and NRRL163 [ATCC1022]) were grown by the Philadelphia VA medical center clinical microbiology lab in 40 ml BACTEC Myco/F Lytic Culture Vials (BD, Sparks, MD) at 30 °C for ten days. Conditioned medium (CM) was then extracted and filtered sequentially through 0.45 µm and 0.2 µm filters as previously described.[3] Aliquots of conditioned media were stored at -80 °C and thawed immediately before use.

**Generation of primary human sinonasal ALI cultures (ALIs) from residual surgical**

Human sinonasal epithelial cells were enzymatically dissociated and grown to confluence in 50% DMEM/Ham's F-12 plus 50% bronchial epithelial basal media (BEBM, Lonza) for 7 days.[1, 2, 4] Dissociated cells were then seeded on Transwell filters (Corning) coated with type I bovine collagen, fibronectin, and bovine serum albumin. Culture medium was removed from the upper compartment after 5-7 days, and cells were fed basolaterally with differentiation medium containing 50% DMEM and 50% BEBM plus Lonza B-ALI Singlequot hEGF (0.5 ng/ ml), epinephrine (5 ng/ml), BPE (0.13 mg/ml), hydrocortisone (0.5 ng/ml), insulin (5 ng/ml), triiodothyronine (6.5 ng/ml), and transferrin (0.5 ng/ml), supplemented with 100 U/ml penicillin, 100 µg/ml streptomycin, 0.1 nM retinoic acid (B-ALI inducer; added fresh for each feeding) as described. For ELISA measurements of ALI culture lysate, a lysis buffer containing no SDS was used, containing 150 mM NaCl, 1 mM EDTA, 0.3% Triton X-100, 0.3% Tween-20, and 100 mM Tris, pH 7.4, with Roche Complete Protease Inhibitor Cocktail. All lysates were normalized by addition of excess lysis buffer to a protein concentration of 2 mg/ml, measured using a Bio-Rad DC protein assay.

**Generation of mouse nasal septum ALI cultures**

Nasal septum was removed from C57BL/6J Wt and PAR-2 knockout (B6.Cg-*F2rl1*$^{tm1Mslb}$/J; Jackson Labs; Bar Harbor, ME USA) mice euthanized for other experimental purposes with IACUC approval according to NIH guidelines, the principles of ARRIVE, and the Basel Declaration. No animals were sacrificed solely for the experiments in this study. Nasal septum was dissociated and cultured at air-liquid interface as described[2] and used after 3 weeks exposure to air for full differentiation in DMEM/F12K media containing 2% NuSerum (Corning), 100 U/ml penicillin, 100 µg/ml streptomycin.

**Quantitative (q) PCR**

RNA was isolated from ALI cultures or stripped turbinate or polyp epithelium as previously described[5] and qPCR was performed using TaqMan primers (ThermoFisher Scientific) for F2RL1 (Hs00608346_m1), ACTB (Hs01060665_g1), and GAPDH (Hs02786623_g1) in separate reactions and relative expression was calculated by calculated by means of the 2-ΔΔ Ct method.

#### Measurement of FITC-conjugated dextran permeability

FITC-dextran (10 kDa) in phenol red-free DMEM was placed on the apical side of the culture, and basolateral solution (also phenol red-free DMEM) was collected from the basolateral side after 30 min incubation at 37°C. Samples were read in a Tecan Spark 10M fluorescence plate reader at 485 nm excitation and 525 nm emission. Basolateral media from cells treated with apical media only (not FITC-dextran) was used for background subtraction.

#### Measurement of airway surface liquid glucose

Glucose permeability was measured by adding 30 µL glucose-free, phenol red-free DMEM into the apical compartment of 0.33cm$^2$ transwells with normal differentiation media placed on the basolateral side. After 6 hrs incubation at 37°C to allow glucose to equilibrate, 25 µL apical solution was removed to measure glucose concentration using a colorimetric kit (Cayman Chemical) as indicated by the manufacturer's instructions and as previously described.[6]

#### Measurement of ciliary beat frequency (CBF)

Cultures were imaged on a Leica microscope with 20x 0.8 NA objective and Basler A602f camera running at 100 frames per second. Experiments were carried out at 28°C. ALIs were washed into HEPES-buffered HBSS containing 1.8 mM $Ca^{2+}$.

#### Imaging of ASL height

Cultures were first washed twice with 25 µM DTT (5 min prewash) to remove large clumps of airway mucus, followed by at least 5-7 washes with PBS alone to remove residual DTT. ASL was labeled with Texas red dextran (10,000 MW; 1 mg/ml) sonicated in perfluorocarbon (PFC-77) ± 2FLI (0.5 mg/ml) and placed in a humidified, heated, gassed (5% $CO_2$) stage-top chamber (Tokai Hit; Tokyo, Japan). The apical side of the cultures was otherwise unmodified; any further drug additions were made basolaterally. Spinning disk confocal images were taken using a 60x (1.0 NA) water immersion objective with 0.2 µm step size. ASL height was determined as described[7] using a 70% of maximum fluorescence threshold to define ASL boundaries. For each culture, three separate fields were analyzed, with the averaged value treated as a single data point.

**Imaging of $Ca^{2+}$ in cells and tissue explants**

Submerged RMPI 2650 and NCI-H520 cells were loaded with 5 μM Fluo-4 for 45 min and $Ca^{2+}$ was imaged as described.[1, 2, 4] Primary human ALIs were loaded as described[1, 2, 4, 5] with 10 μM Fluo-4 for 90 min on the apical side. Imaging was performed using a Nikon TS-100 microscope with equipped with 10x 0.3 NA PlanFluor objective (for ALIs) or 20x 0.8 NA PlanApo objective (for submerged cells) and standard FITC/GFP filter set. Mouse ALIs were loaded similarly with Calbryte 590 to avoid green autofluorescence and visualized with TRITC filter set. Background was estimated by imaging unloaded ALIs or off cell region for submerged cells at identical settings as described.[5] Experiments utilized Hank's Balanced Salt Solution (HBSS) buffered with 20 mM HEPES at pH 7.4 with 1.5 mM $Ca^{2+}$ and either 5.5 mM glucose (basolateral side) or 0 glucose (apical side). Experiments performed in the absence of $Ca^{2+}$ (0-$Ca^{2+}$) utilized cultures loaded with BAPTA-AM (10 μM; 15 min preincubation) and extracellular solutions containing no added $Ca^{2+}$ and 1 mM EGTA.

Flat thin sections of mucosa (≥1 mM thick) were cut from middle turbinate or nasal polyp tissue and mounted in an oblong EasyMount Ussing Chamber insert (Harvard Instruments) to isolate the apical and basolateral surfaces. The tissue and insert were then placed apical side up in a petri dish containing HBSS containing 1x MEM Amino Acids, 1x MEM Vitamins, 2 mM L-glutamine. The apical side was loaded with 15 μM Calbryte 590-AM in the HBSS solution as above contaning 0.4% pluronic F127 for 2 hours. Insert was then washed and secured apical side down with vacuum grease on top of a perfusion chamber (Warner Instruments, RC-26GLP) with a low channel and glass coverslip bottom held on a heated stage (Warner PH-1) and perfused with 37°C Tyrode's buffer, containing (in mM): 140 NaCl, 5 KCl, 2 $CaCl_2$, 1 $MgCl_2$, 10 glucose, 5 NaPyruvate, 10 HEPES, pH 7.4. Imaging was performed on an inverted microscope (Olympus IX83) with a standard TRITC filter set using a 10x 0.3 NA objective and spinning disk confocal unit (Olympus DSU), XCite 120 LED Boost light source (Excelitas Technologies), Orca Flash 4.0 sCMOS camera (Hammamatsu), and Metafluor using 250 ms exposure and 4x4 binning and 5 sec sampling frequency to minimize dye bleaching due to the high excitation intensity needed due to the spinning disk.

**Table S1:** Clinical characteristics of patients from whom tissue was used.

**Non-CF patients from whom tissue was used for ALI culture**

| Patient | Age at Surgery | Gender | Ethnicity | Diagnosis | # Prior FESS | Polyps | Lund-Mackay | SNOT-22 | Smoking History | Asthma | AFS | Abx | Steroids | Comorbidities |
|---|---|---|---|---|---|---|---|---|---|---|---|---|---|---|
| 1 | 63 | Female | African American | CRS | 0 | No | N/A | N/A | Yes | No | No | No | No | HTN, DM |
| 2 | 48 | Female | Caucasian | CRS | 2 | No | N/A | 49 | No | No | No | No | No | Allergies |
| 3 | 51 | Female | Caucasian | CRS | 0 | No | 10 | 64 | Yes | No | No | No | Yes | Depression |
| 4 | 51 | Male | Caucasian | CRS | 1 | Yes | 9 | 62 | Yes | Yes | No | Yes | Yes | Allergies, GERD |
| 5 | 62 | Female | Caucasian | CRS | 0 | No | 4 | 53 | No | Yes | No | Yes | Yes | Allergies, HTN, OSA |
| 6 | 67 | Female | Caucasian | CRS | 0 | Yes | 14 | 20 | No | Yes | Yes | No | No | GERD |
| 7 | 70 | Male | Caucasian | CRS | 0 | No | 11 | 39 | No | No | No | No | No | DM, GERD |
| 8 | 44 | Female | Caucasian | CRS | 0 | No | 6 | 27 | No | Yes | No | No | Yes | Pacemaker |
| 9 | 52 | Male | Caucasian | Fungal Ball | 0 | Yes | 15 | 35 | No | No | Yes | No | No | Allergies |
| 10 | 60 | Female | Caucasian | CRS | 2 | Yes | 16 | 53 | No | Yes | No | No | No | Samter's Triad, Allergies |
| 11 | 61 | Female | Caucasian | CRS | 0 | Yes | N/A | 46 | Yes | Yes | No | Yes | Yes | HTN |
| 12 | 37 | Male | Caucasian | CRS | 1 | Yes | 18 | 68 | Yes | No | No | No | No | Allergies, Sinonasal Trauma |
| 13 | 24 | Female | Caucasian | CRS | 0 | No | 13 | 69 | No | No | No | No | No | Allergies, GERD |
| 14 | 83 | Male | Caucasian | CRS | 1 | Yes | 3 | 10 | No | No | No | No | No | HTN |
| 15 | 57 | Male | Caucasian | CRS | 0 | Yes | 15 | 16 | No | No | No | Yes | No | Allergies |
| 16 | 19 | Male | Caucasian | CRS | 1 | Yes | N/A | 41 | No | Yes | No | No | No | Allergies |
| 17 | 49 | Male | Caucasian | CRS | 2 | Yes | 12 | 11 | No | No | No | No | No | Pulmonic Stenosis, Allergies |
| 18 | 70 | Female | Caucasian | CRS | 0 | No | 8 | 61 | No | No | No | Yes | Yes | HTN |
| 19 | 34 | Male | Caucasian | CRS | 0 | Yes | N/A | N/A | No | No | No | No | No | Allergies |
| 20 | 71 | Male | Caucasian | CRS | 2 | Yes | 24 | 8 | No | Yes | No | No | Yes | Samters, Hypothyroid |
| 21 | 71 | Male | Caucasian | CRS | 4 | Yes | 18 | 13 | No | No | No | No | No | Samter's triad, Allergies, GERD, AERD, OSA, HTN |
| 22 | 60 | Male | Caucasian | CRS | 1 | Yes | 11 | 57 | No | No | No | No | Yes | N/A |
| 23 | 59 | Female | Caucasian | CRS | 0 | No | N/A | 61 | No | No | No | No | No | DM, GERD, HTN |
| 24 | 66 | Male | Caucasian | CRS | 0 | Yes | N/A | 18 | No | No | No | No | Yes | N/A |
| 25 | 60 | Male | Caucasian | CRS | 0 | Yes | N/A | N/A | No | Yes | No | No | No | N/A |
| 26 | 46 | Female | Hispanic, Latino | CRS | 1 | Yes | N/A | 95 | No | Yes | No | No | Yes | Obesity |
| 27 | 48 | Male | Caucasian | CRS | 0 | No | N/A | N/A | No | No | No | No | Yes | OSA, COPD |
| 28 | 56 | Female | Caucasian | CRS | 0 | Yes | N/A | 27 | No | No | No | No | No | N/A |
| 29 | 69 | Male | Caucasian | CRS | 3 | Yes | N/A | 12 | No | No | No | No | No | N/A |
| 30 | 44 | Male | Caucasian | CRS | 2 | Yes | 7 | 22 | No | No | No | Yes | No | Allergies, HTN |
| 31 | 44 | Female | Caucasian | CRS | 0 | Yes | 20 | 30 | No | Yes | No | No | No | HTN, Diabetes |
| 32 | 64 | Male | Caucasian | CRS | 3+ | Yes | 20 | 32 | No | No | No | No | Yes | Samter's triad, Allergies |
| 33 | 32 | Male | Caucasian | CRS | 1 | Yes | 10 | 42 | No | No | No | No | Yes | Allergies, GERD |
| 34 | 52 | Male | Caucasian | CRS | 2 | No | 16 | 11 | No | No | No | No | Yes | DM, GERD |
| 35 | 75 | Male | Caucasian | CRS | 2 | No | 12 | 68 | No | Yes | No | No | No | Allergies, GERD |

**Patients from whom tissue was used for turbinate tissue calcium imaging and ALI culture**

| Patient | Age at Surgery | Gender | Ethnicity | Diagnosis | # Prior FESS | Polyps | Lund-Mackay | SNOT-22 | Smoking History | Asthma | AFS | Abx | Steroids | Comorbidities |
|---|---|---|---|---|---|---|---|---|---|---|---|---|---|---|
| 1 | 60 | Female | Caucasian | inverted papilloma | 0 | No | 14 | 9 | Yes | Yes | No | No | Yes | DM, HTN, OSA |
| 2 | 66 | Female | Caucasian | skull base tumor | 4 | No | N/A | 70 | Yes | No | No | No | No | ARS, DM, HHT |
| 3 | 78 | Female | African American | skull base tumor | 0 | No | N/A | 19 | No | No | No | No | No | HTN, Allergies |
| 4 | 56 | Female | Caucasian | CSF leak | 0 | No | N/A | N/A | No | No | No | No | No | Hypothyroid |
| 5 | 76 | Female | Caucasian | Pituitary microadenoma | 0 | No | N/A | N/A | No | No | No | No | No | HTN, CAD, PVD |
| 6 | 21 | Male | Caucasian | Orbital decompression | 0 | No | 0 | 0 | Yes | No | No | No | No | N/A |
| 7 | 65 | Male | Caucasian | inverted papilloma | 0 | No | 4 | N/A | No | No | No | No | Yes | N/A |

**Patients from whom tissue was used for polyp tissue calcium imaging and ALI culture**

| Patient | Age at Surgery | Gender | Ethnicity | Diagnosis | # Prior FESS | Polyps | Lund-Mackay | SNOT-22 | Smoking History | Asthma | AFS | Abx | Steroids | Comorbidities |
|---|---|---|---|---|---|---|---|---|---|---|---|---|---|---|
| 1 | 57 | Male | Caucasian | CRS | 1 | Yes | 10 | 53 | No | No | No | No | Yes | GERD |
| 2 | 38 | Male | Caucasian | CRS | 1 | Yes | 20 | 67 | No | Yes | No | No | Yes | N/A |
| 3 | 66 | Male | Caucasian | CRS | 0 | Yes | 13 | 22 | No | No | No | No | Yes | HTN |
| 4 | 56 | Male | Caucasian | CRS | 1 | Yes | 16 | 41 | No | No | No | No | Yes | HTN, OSA |
| 5 | 34 | Female | Caucasian | CRS | 0 | Yes | 6 | 34 | No | Yes | No | No | No | Allergies |
| 6 | 43 | Male | Caucasian | CRS | 2 | Yes | 20 | 58 | No | No | Yes | Yes | Yes | Samter's triad, Allergies, HTN |
| 7 | 53 | Female | Caucasian | CRS | 1 | Yes | 15 | 14 | No | No | No | No | Yes | Sampter's Triad, Allergies, HTN, DM |

**ΔF508/ΔF508 CF patients from whom tissue was used for ALI culture**

| Patient | Age at Surgery | Gender | Ethnicity | Diagnosis | # Prior FESS | Polyps | Lund-Mackay | SNOT-22 | Smoking History | Asthma | AFS | Abx | Steroids | Comorbidities |
|---|---|---|---|---|---|---|---|---|---|---|---|---|---|---|
| CF1 | 26 | Female | Caucasian | CF-related CRS | 1 | No | N/A | 78 | No | Yes | No | No | No | CF, GERD, DM |
| CF2 | 23 | Male | Caucasian | CF-related CRS | 1 | Yes | N/A | N/A | No | No | No | No | No | CF, GERD |
| CF3 | 33 | Female | Caucasian | CF-related CRS | 1 | Yes | 13 | 8 | No | No | No | No | No | CF, Double Lung Transplant, Allergies, GERD, HTN, DM |
| CF4 | 32 | Female | Caucasian | CF-related CRS | 1 | Yes | N/A | 77 | No | Yes | No | No | No | CF, GERD |
| CF5 | 58 | Female | Caucasian | CF-related CRS | 2 | Yes | 18 | 27 | No | Yes | No | No | No | CF, Allergies, GERD, HTN |

Parameters of patients from whom primary cells were grown and/or tissue was used for experiments in the main text, including Lund-Mackay[8, 9] and SNOT-22 scores.[10] Patients with a history of systemic inheritable disease (eg, granulomatosis with polyangiitis, systemic immunodeficiencies) were excluded, as well as patients receiving antibiotics, oral corticosteroids, or anti-biologics (e.g. Xolair) within one month of surgery. Previously, cultures from control and CRS patients as well as cultures derived from turbinate vs ethmoid sinus were compared for CBF and calcium responses to PAR-2 stimulation, and no significant differences were observed.[2] As previously described,[2, 3, 5-7, 11-14] we find that once cells are removed from an inflammatory environment and expanded and cultured for 3-6 weeks to differentiated ALIs in defined media, secondary disease-related phenotypes are removed and cells reflect a "healthy" baseline state, with responses overwhelmingly dictated by genetics. This allows disease-relevant *in vitro* manipulations (IL-13, CSC, etc.) with comparison of unmanipulated cells from the same patient as "control."

**Abbreviations:** Abx, history of antibiotics; AFS, allergic fungal sinusitis; ARS, allergic rhinosinusitis; CRS, chronic rhinosinusitis; DM, diabetes mellitus; FESS, functional endoscopic sinus surgery; GERD, gastroesophageal reflux disease; HHT, heterditary hemorrhagic telangiectasia; HTN, hypertension; IP, inverted papilloma; N/A, not available; OSA, obstructive sleep apnea; SNOT-22, 22 question sinonasal outcomes test;

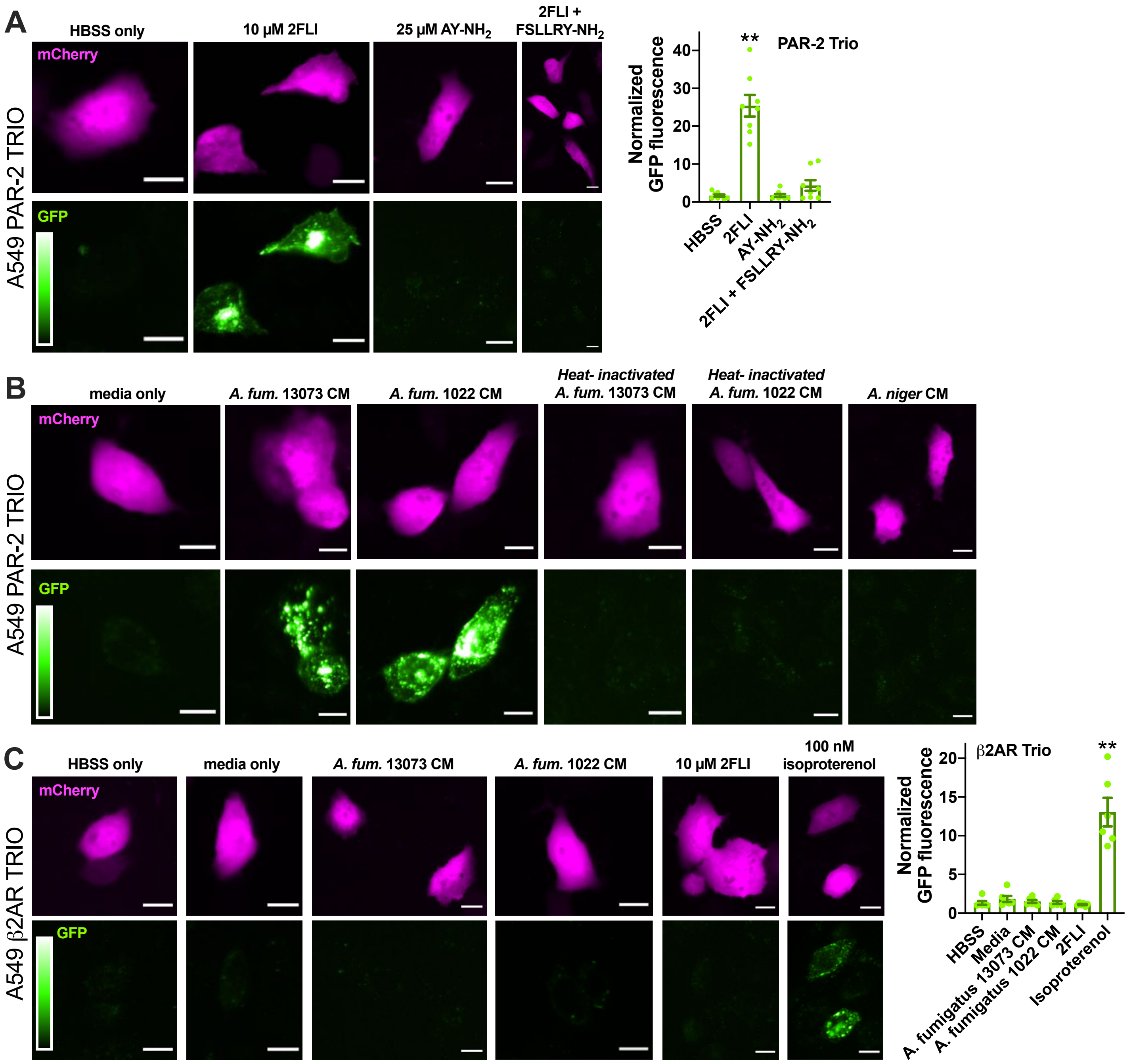


**FIG S1. Activation of PAR-2 by peptide Par-2 agonist and *Aspergillus fumigatus* conditioned media (CM) in A549 cells.** Cells were transfected with the PAR-2 Trio system[15] as described in the main text to detect receptor-β-arrestin interaction upon PAR-2 activation. Many other studies have already demonstrated that β-arrestin is recruited to PAR-2 upon activation.[15-22] Using this assay allows to directly look at PAR-2 activation in the absence of any other signals downstream of other targets of proteases and/or CM that could impinge on the PAR-2 signaling pathway. **A,** After 90 min stimulation with PAR-2 agonist 2-furoyl-LIGRLO-amide (2FLI), GFP fluorescence was increased ~25 fold compared with buffer (Hank's balanced salt solution; HBSS) only or PAR-4 activating peptide AY-$NH_2$. The GFP fluorescence increase, signaling PAR-2 activation, was lost when cells were stimulated with 2FLI in the presence of 10 μM PAR-2 antagonist FSLLRY-$NH_2$. **B,** Representative images of PAR-2 activation with *A. fumigatus* CM as quantified in main text Figure 1E. Cells were exposed to conditioned media for 10 min followed by washing and resuspension in HBSS for 90 min. **C,** A549 cells were transfected with β2 adrenergic receptor (β2AR) Trio. *A. fumigatus* CM did not activate β2AR while 100 nM isoproterenol did. Bar graph shows quantification of results. All data points in A and C are from ≥7 independent experiments imaging fields from independent transfected wells on separate days. Significance was determined by 1-way ANOVA with Dunnett's posttest comparing all values to HBSS alone; $**p<0.01$.

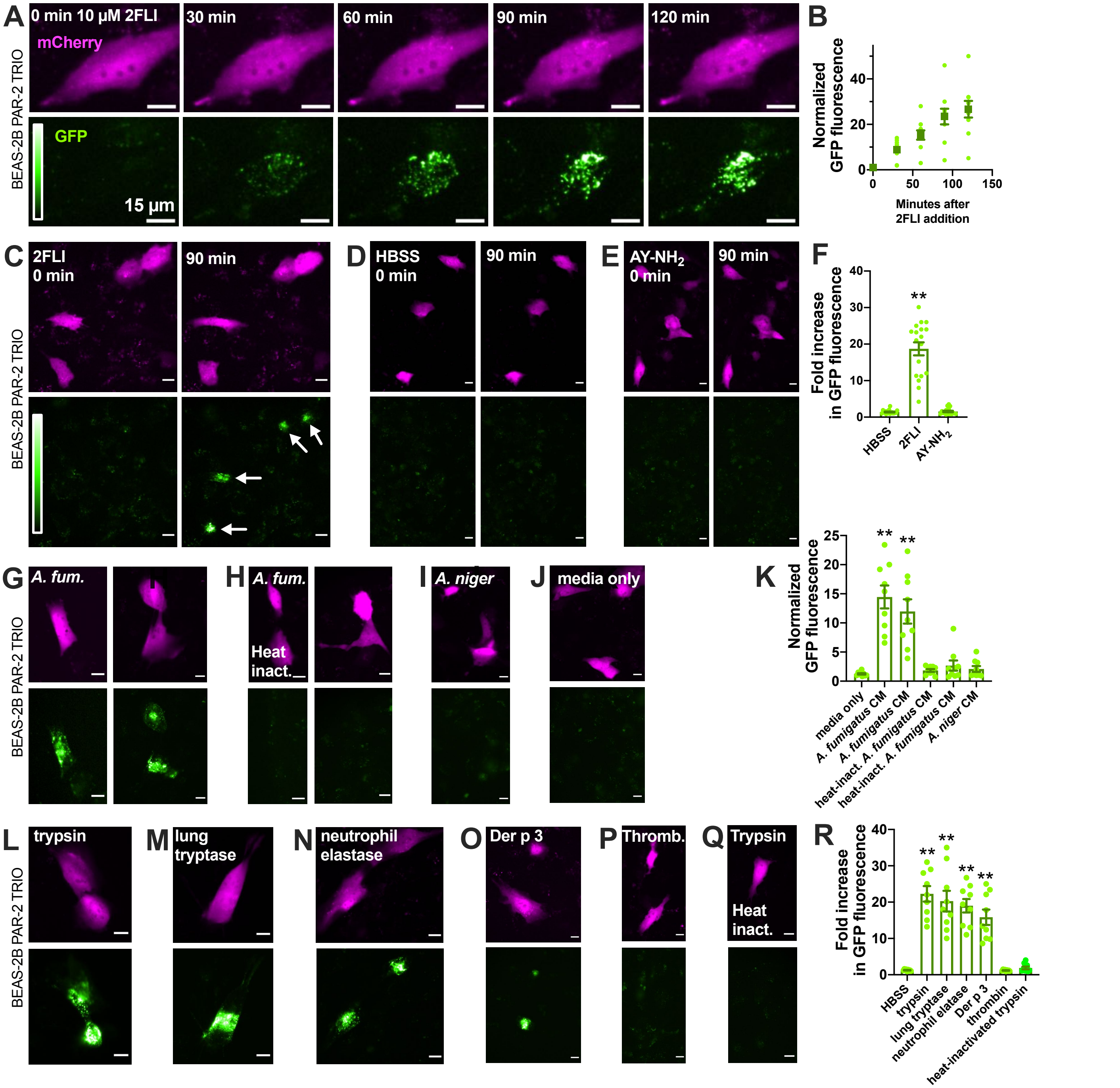


**FIG S2.** ***Aspergillus fumigatus*** **CM and proteases activate PAR-2 in Beas-2B immortalized squamous bronchial epithelial cells. A-B,** Representative images (*A*) and quantification (*B*) showing time-course of PAR-2-arrestin recruitment and resulting increase in GFP fluorescence in Beas-2B cells transfected with the PAR-2 Trio assay components and stimulated with PAR-2 agonist 2FLI.[15] **C-F,** After 90 min stimulation, 2FLI (*C*) but not HBSS alone (*D*) or AY-NH2 (*E*) resulted in ~20-fold increased GFP fluorescence, quantified in *F*. **G-K,** Cells were stimulated for 10 min with 25% *A. fumigatus* CM (*G*, strain 13073 left and 1022 right), heat-inactivated *A. fumigatus* CM (*H*; 20 min; 100 °C; strain 13073 left and 1022 right), A. niger CM (*I*), or media only (*J*) diluted in HBSS, followed by washing with HBSS and incubation for 90 min. GFP quantification is shown in *K*. **L-R,** Cells were stimulated for 10 min with 25 nM trypsin (L), human lung tryptase (M), neutrophil elastase (N), Der p 3 (O), thrombin (P), or heat-inactivated trypsin (Q), followed by washing with HBSS and incubation for 90 min. Quantification of GFP fluorescence shown in *R*. Data points in *B*, *F*, *K*, and *R* are independent experiments done on different days (n ≥5). Significance was determined by 1-way ANOVA with Dunnett’s posttest comparing all values to HBSS alone; ***p*<0.01.

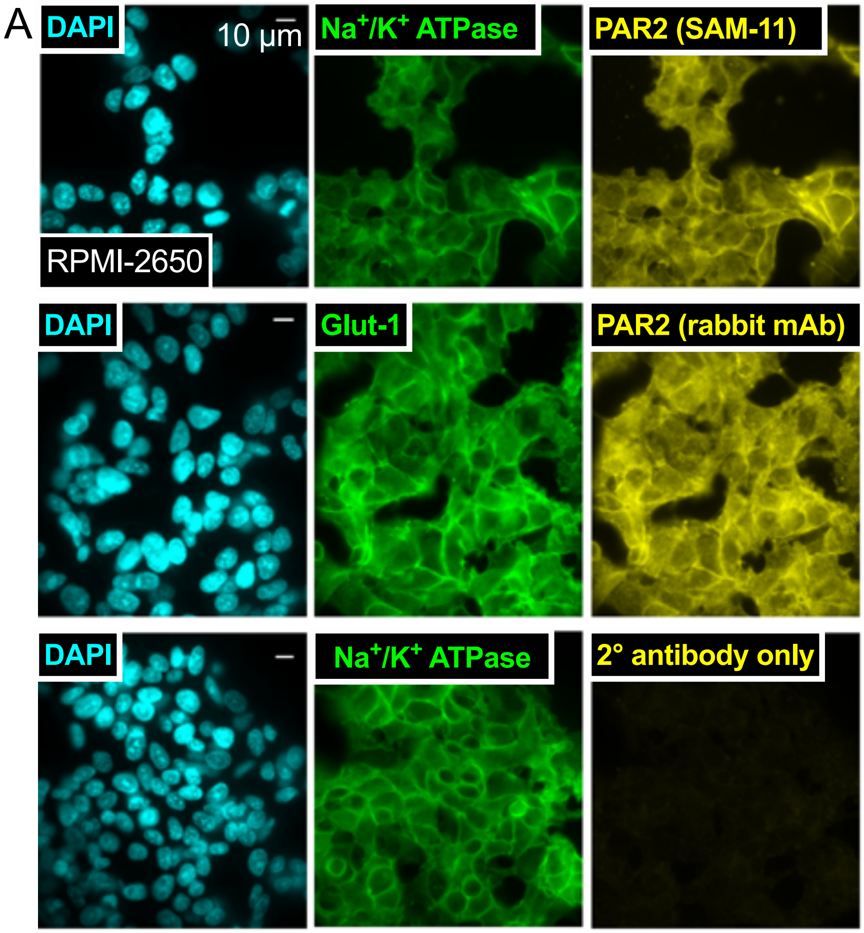


**FIG S3. PAR-2 expression in RPMI 2650 cells.** We examined nasal septal squamous cell line RPMI 2650,[23] which has a near diploid karyotype[24, 25] and forms tight junctions (~150-200 Ω•cm$^2$) when cultured at air-liquid interface (ALI).[26-29] Immunofluorescence with two different primary antibodies revealed plasma membrane localization of PAR-2 in RPMI 2650 cells, suggested by similar staining patterns with $Na^+/K^+$ ATPase and Glut1. Representative results shown from 3 independent experiments conducted on separate days.

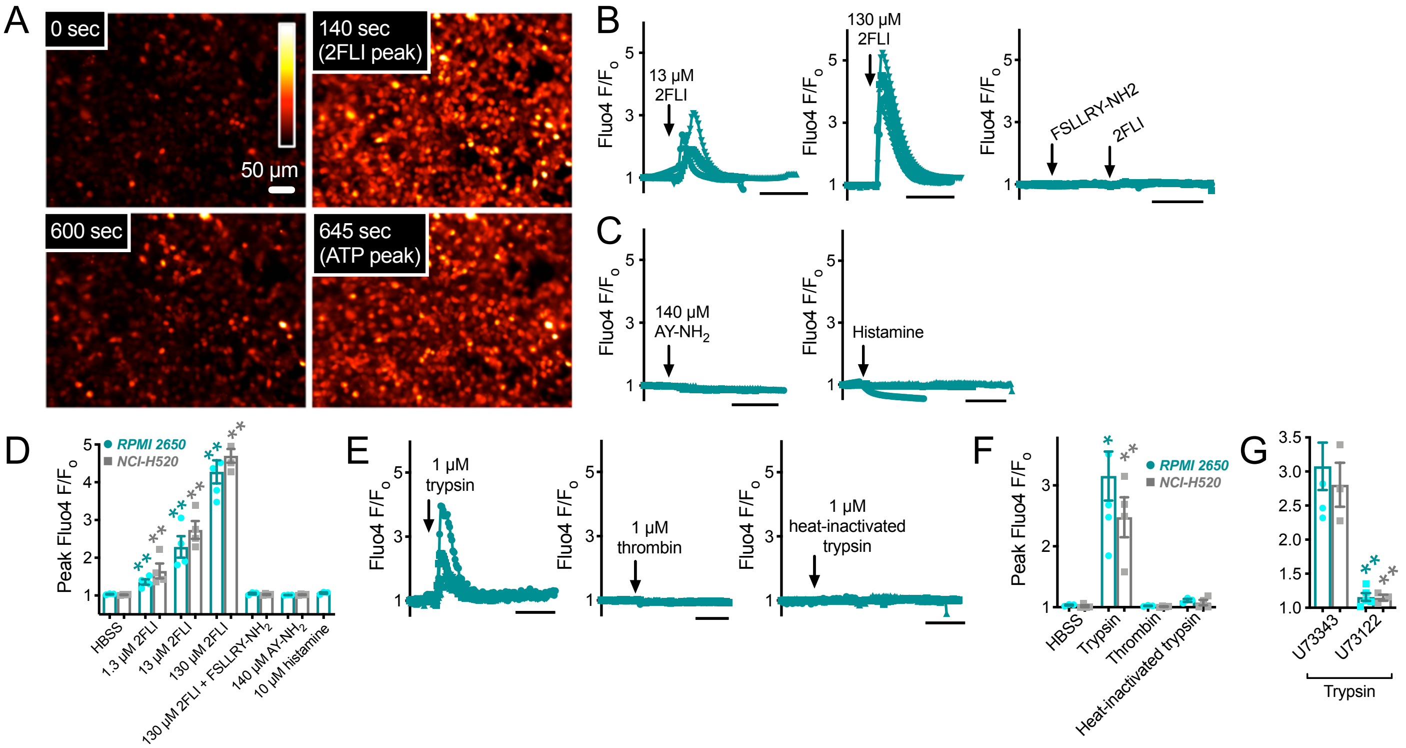


**FIG S4. PAR-2 function in RPMI 2650 and NCI-H520 squamous airway cell lines. A,** We imaged intracellular $Ca^{2+}$ ($[Ca^{2+}]_i$) dynamics using the fluo-4. In submerged sub-confluent RPMI 2650 cells, we observed a $[Ca^{2+}]_i$ transient in response to stimulation with PAR-2 agonist 2-furoyl-LIGRLO-amide (2FLI). Single experiment shown, representative of 3 independent experiments. Representative images are pseudocolored and scaled linearly to show fluo-4 fluorescence changes. ATP is used as a positive control as most airway cells we gave encountered express purinergic receptors. **B,** 2FLI dose-dependently increased $[Ca^{2+}]_i$. Plotted are individual traces from 3-4 separate experiments. **C,** An agonist for PAR-4 (AY-$NH_2$) and histamine did not affect intracellular $Ca^{2+}$. Plotted are individual traces from 3-4 separate experiments. **D,** Peak fluo-4 data are shown from experiments as in *B-C*. Similar results were observed in NCI-H520 cells. Significance determined by 1-way ANOVA with Dunnett's posttest comparing values to HBSS alone (control); ** *p* <0.01. **E,** Supporting expression of PAR-2 but not PAR-4, we found that trypsin, which activates PAR-2 and PAR-4, stimulated a $[Ca^{2+}]_i$ response while thrombin, which activates only PAR-4, did not. Heat inactivated trypsin (20 min, 90°C) did not cause a $[Ca^{2+}]_i$ increase. Plotted are individual traces from 3-4 separate experiments. **F,** Peak fluo-4 data for trypsin and thrombin are summarized from experiments in *E*. Significance determined by 1-way ANOVA with Dunnett's posttest comparing values to HBSS alone (control); *$p$ <0.05 and **$p$ <0.01 **G,** The trypsin-activated $[Ca^{2+}]_i$ response was blocked by the phospholipase C (PLC) inhibitor U73122 (1 µM; 20 min pre-treatment) but not by its inactive analogue U73343. Peak fluo-4 values are plotted from 3-5 independent experiments. Significance determined by 1-way ANOVA with Bonferroni posttest for paired comparisons.

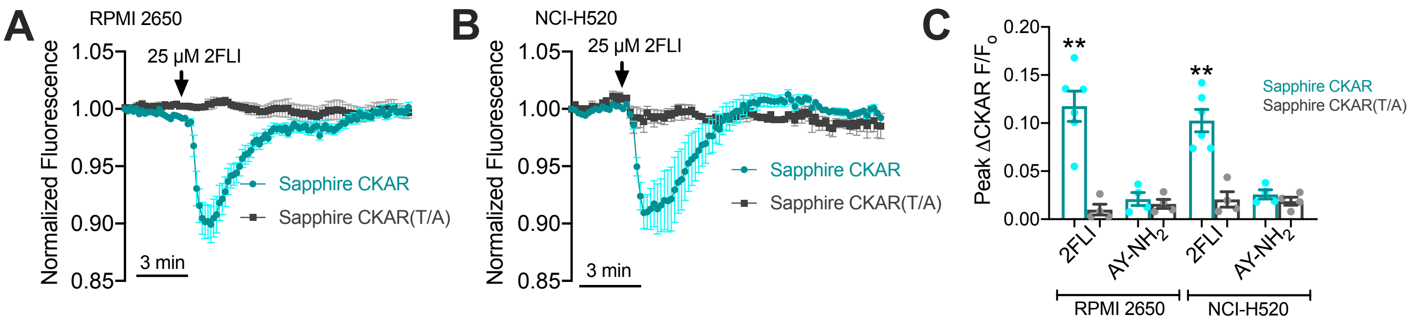


**FIG S5. Activation of protein kinase C (PKC) during PAR-2 stimulation in airway squamous cells.** Protein kinase C has been implicated in chronic inflammatory diseases.[3, 30, 31] We examined protein kinase C signaling in RPMI 2650 and NCI-H520 cells expressing a fluorescent PKC biosensor, Sapphire CKAR, which decreases in fluorescence with PKC phosphorylation.[32] **A-B)** Stimulation with PAR-2 agonist 2FLI caused a transient decrease in Sapphire CKAR fluorescence (signaling PKC activation) in RPMI 2650 cells (*A*) and NCI-H520 cells (*B*) that was not observed with a mutated form of the sensor that cannot be phosphorylated by PKC, Sapphire CKAR(T/A). **C)** Quantification of independent experiments (n = 5) showing activation of PKC signaling (change in Sapphire CKAR fluorescence compared with Sapphire CKAR(T/A) with Par-2 agonist 2FLI but not Par-4 agonist AY-$NH_2$. Significance determined by 1-way ANOVA with Bonferroni posttest comparing Sapphire CKAR with Sapphire CKAR(T/A) for each condition.

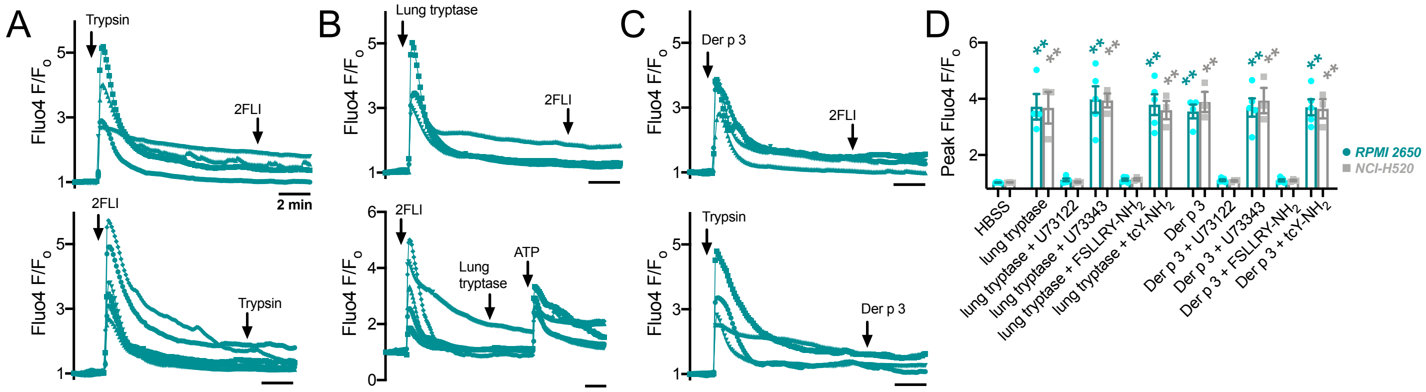


**FIG S6. Activation of PAR-2 by endogenous and exogenous proteases in airway squamous cells. A)** When 2FLI (25 µM) was applied to RPMI 2650 cells after trypsin (100 nM) or vice versa, no further response was observed, suggesting the 2FLI add trypsin stimulate $Ca^{2+}$ through the same pathway, likely PAR-2. **B)** A similar phenotype was observed when 2FLI was applied after human lung tryptase (50 nM) or when tryptase was applied after 2FLI, suggesting tryptase activates PAR-2. Purinergic agonist ATP did activate a response when applied after 2FLI and tryptase. **C)** We also tested the dust mite protease allergen Der p 3 (50 nM), and find that 2FLI did not evoke a response after Der p 3 application, and Der p 3 did not activate a response after trypsin application. Plotted on graphs in *A-C* are traces from 3-5 individual experiments. **D)** Peak fluo-4 values are shown from experiments as in *A-C*. $[Ca^{2+}]_i$ responses to tryptase or Der p 3 were blocked in RPMI 2650 and NCI-H520 cells by U73122 but not U73343 (10 µM, 30 min pretreatment), supporting PLC activation by a GPCR. The responses were also inhibited by a PAR-2 antagonist (FSLLRY-$NH_2$; 50 µM) but not a PAR-4 antagonist (tcY-$NH_2$; 50 µM). The above data support functional PAR-2, but not PAR-4, expression in airway squamous cells and coupling of PAR-2 to $[Ca^{2+}]_i$. Significance determined by 1-way ANOVA with Dunnett’s posttest comparing all values to control (HBSS only); ** *p* <0.01.

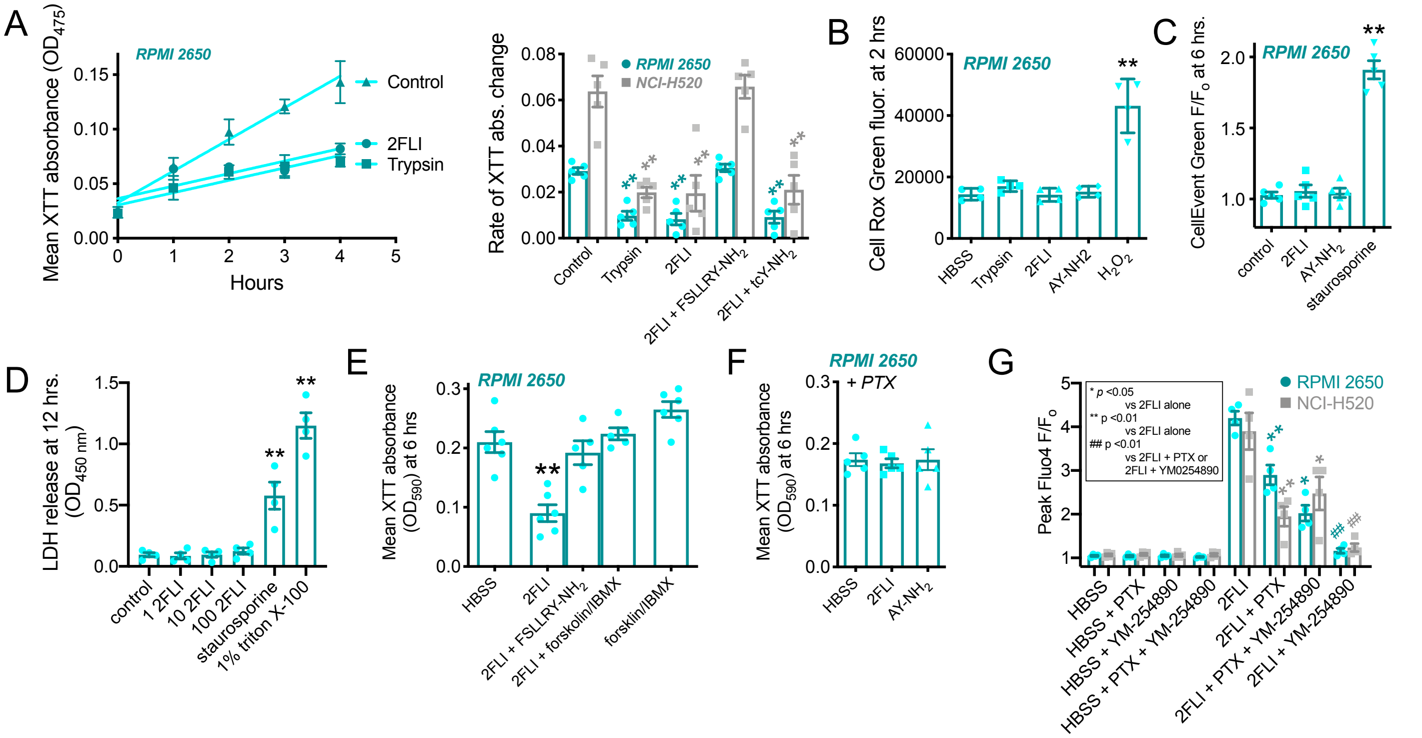


**FIG S7. Effects of PAR-2 stimulation on airway squamous cell metabolism. A,** PAR-2 has been suggested to be both a suppressor as well as a promotor or tumor growth.[33, 34] Surprisingly, we found that slowed metabolism of RPMI 2650 cells. PAR-2 activation with trypsin or 2FLI reduced metabolism over 4 hours in RPMI 2650 and NCI-H520 cells, as measured by XTT assay, which is an indirect readout of metabolism via NADH production. This was blocked by a PAR-2 antagonist FSLLRY-$NH_2$ but not PAR-4 antagonist tcY-$NH_2$. Real time changes in XTT absorbance shown on left, and rate of change (linear fit; abs/hr) plotted on right. Significance determined by 1-way ANOVA with Dunnett's posttest comparing values to control (media only); ** $p$ <0.01. **B,** PAR-2 activation did not not increase oxidative stress, as measured by CellRox Green fluorescence. 0.1% $H_2O_2$ used as positive control. Significance determined by 1-way ANOVA with Dunnett's posttest comparing values to control (media only); ** $p$ <0.01. **C,** PAR-2 did not activate Caspases 3 or 7 over 6 hours, as measured by a fluorescent DEVD peptide assays (CellEvent). Staurosporine (PKC inhibitor) used as positive control for activation of apoptosis. Significance determined by 1-way ANOVA with Dunnett's posttest comparing values to control (media only); **$p$ <0.01. **D,** PAR-2 did not activate LDH release over 12 hours. Staurosporine used as positive control as well as cell lysis by 1% triton X-100 detergent. Significance determined by 1-way ANOVA with Dunnett's posttest comparing values to control (media only); ** $p$ <0.01. **E-F,** While XTT assays are sometimes incorrectly used as estimates of proliferation or viability, while really they measure metabolic steady-state NADH levels. Thus, our data support a reduction in metabolic activity during acute PAR-2 activation but not a decrease in viability or cell death, as there was no LDH release. The effects of 2FLI on NADH production were eliminated when cells were treated simultaneously with adenylyl cyclase-activating forskolin and phosphodiesterase inhibitor isobutylmethylxanthine (IBMX) (*E*) or when cells were pretreated with 18 hours with pertussis toxin (PTX; 18 hrs. pretreatment; *F*), which ADP-ribosylates and inactivates G$\alpha_i$ proteins. These data suggest that slowed metabolism and/or proliferation activated by PAR-2 stimulation requires coupling of PAR-2 to G$\alpha_i$ and lowering of intracellular cAMP. Significance determined by 1-way ANOVA with Dunnett's posttest comparing values to control (media only); ** $p$ <0.01. ***G*,** PAR-2 has been previously reported to couple to both G$\alpha_i$ and G$\alpha_q$ in various cell types [35]. While G$\alpha_q$ activates PLC, G$\alpha_i$-linked receptors can also activate $Ca^{2+}$ signals through the G$\beta\gamma$ subunits. Peak Fluo-4 responses in RPMI 2650 and NCI-H520 cells were inhibited ~50% by PTX and ~50% by G$\alpha_q$11 inhibitor YM-254890,[36] while a combination of both inhibitors almost completely blocked $[Ca^{2+}]_i$ responses. These data support coupling of PAR-2 to both G$\alpha_i$ and G$\alpha_q$ in squamous airway epithelial cells. Significance determined by 1-way ANOVA with Bonferroni posttest comparing conditions with 2FLI + inhibitors to 2FLI alone; ** $p$ <0.01.

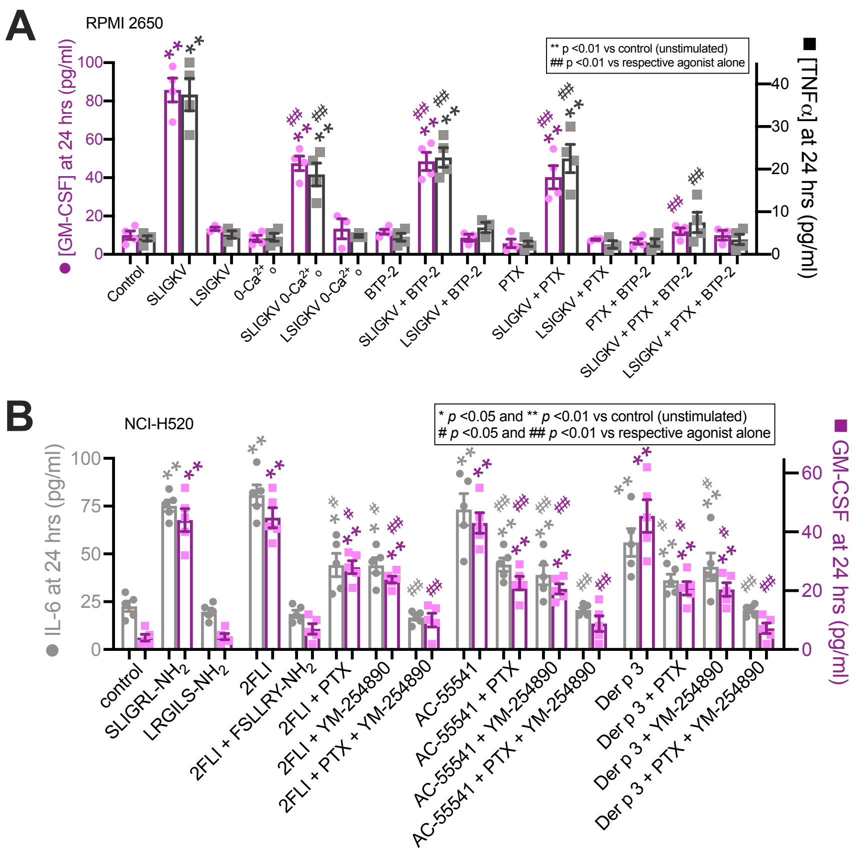


**FIG S8. Effects of PAR-2 stimulation on airway squamous cell inflammatory responses. A,** Squamous epithelial cells can be a source of cytokines and PAR-2 can be an inflammatory receptor.[33, 37] RPMI 2650 cells have been shown to produce cytokines like TGFβ in response to insults like particulate matter, viral infection, and TNFα,[23, 37-39] while H520 cells can produce IL-6 and GM-CSF,[40-42] an important cytokine in allergy and asthma.[43-47] In RPMI 2650 cells, PAR-2 agonist peptide SLIGKV, but not scrambled LSIGKV peptide, increased secretion of TNFα and GM-CSF into the media over 24 hrs as measured by ELISA. Previously, $Ca^{2+}$ influx via store-operated Orai/Stim channels was implicated in PAR-2 induced inflammatory responses.[48] SLIGKV-induced cytokine secretion was reduced ~50% in the absence of extracellular $Ca^{2+}$ ($0\text{-}Ca^{2+}_o$; no added $Ca^{2+}$ plus 2 mM EGTA) as well as in the presence of the store-operated $Ca^{2+}$ channel inhibitor BTP-2 or PTX. A combination of BTP-2 and PTX completely abrogated TNFα and GM-CSF secretion, suggesting that $Ca^{2+}$ influx contributes to activation of cytokine secretion in RPMI 2650 cells, but $G_i$ coupled PAR-2 receptors can still activate inflammation in the absence of $Ca^{2+}$ influx through other pathways (e.g., ERK signaling). **B,** In NCI-H520 cells, PAR-2 agonist peptide SLIGRL-$NH_2$ but not scrambled LRGILS-$NH_2$ increased IL-6 and GM-CSF secretion, and this was mimicked by 2FLI, non-peptide PAR-2 agonist AC-55541, and dust mite protease Der p 3. Responses to 2FLI, AC-55541, and Der p 3 were reduced somewhat by $G\alpha_i$ inhibitor PTX and $G\alpha_q$ inhibitor YM-254890, but complete inhibition was only observed with both compounds, again supporting a dual coupling of PAR-2 to $G\alpha_i$ and $G\alpha_q$ in squamous airway epithelial cells. All points are individual experiments (n = 3-6). Significances determined by 1-way ANOVA with Bonferroni posttest for paired comparisons.

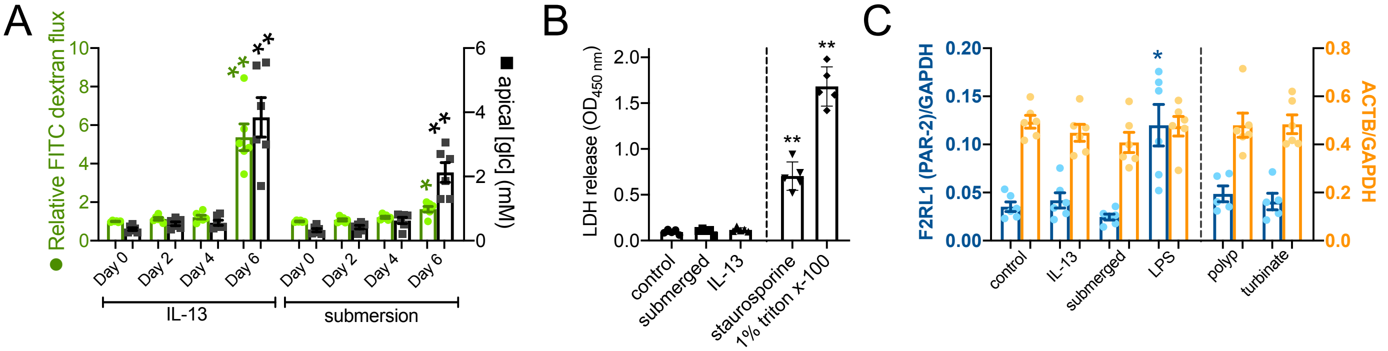


**FIG S9. Epithelial barrier permeability, LDH release, and PAR-2 transcript levels with IL-13 treatment or submersion in primary nasal ALIs. A,** FITC dextran permeability (green), which will increase with epithelial barrier dysfunction, and apical glucose concentration (black), which will increase with epithelial barrier dysfunction, were measured with IL-13 and submersion treatment as indicated. Each data point is an independent ALI from a different patient (n = 5 total ALIs). Significance at each time point by 1-way ANOVA with Bonferronni posttest; ***p*<0.01. **B,** LDH release into cell culture media was measured via colormetric assay as described in the text. Staurosporine (100 ng/ml; 4 hrs) and 1% triton X-100 were used as controls to induce apoptotic death and nonspecific lysis, respectively. Each data point is an independent ALI from a different patient (n = 5 total ALIs). Significance by 1-way ANOVA with Dunnett posttest; ***p*<0.01. **C,** Transcript levels of PAR-2 (F2RL1 gene, relative to GAPDH) in ALIs after 25 days (control), 21 days + 4 days of IL-13, 21 days + 4 subsequent days of submersion, or 23 days + 2 days of LPS (20 μg/ml). Also shown are polyp and turbinate epithelium samples (n = 5 samples from 5 individual patients). Actin (ACTB) shown as a control. Significance determined by 1-way ANOVA with Bonferronni post test; * *p*<0.05.

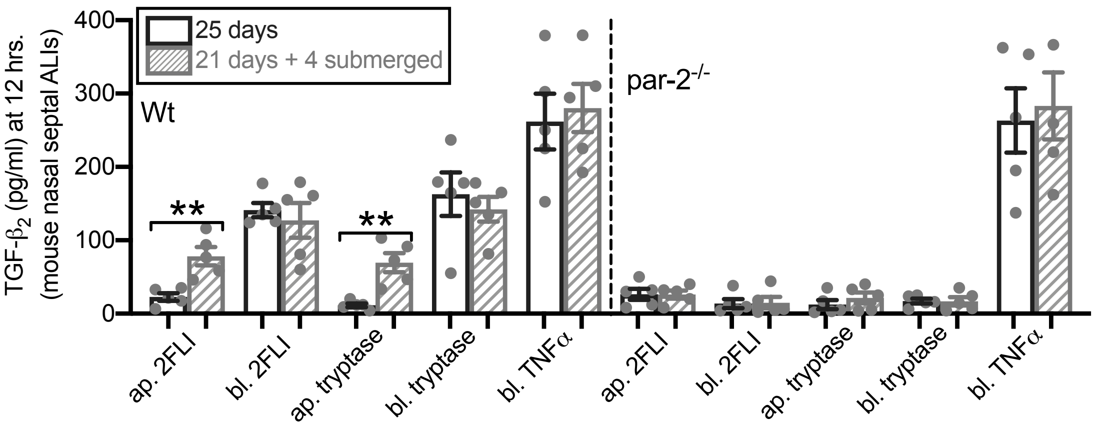


**FIG S10. Differential apical vs basolateral 2FLI responses with submersion are dependent on PAR-2.** Mouse nasal ALIs cultured from Wt or PAR-2 knockout (par-2$^{-/-}$) mouse tissue were grown stimulated with apical vs basolateral 2FLI or tryptase or basolateral TNFα. Wt ALIs, but not par2$^{-/-}$ ALIs, exhibited responses to basolateral 2FLI or tryptase, while only Wt ALIs exposed to submersion to induce squamous differentiation responded to apical 2FLI or tryptase. Both Wt and par2-/- ALIs responded to TNFα. Each point is one single ALI from independent an experiment (4-6 per condition). Each experiment using primary human cells (B-C) used cells from a different individual patients. Each condition with primary mouse cells used cells grown from 2-3 different mice. Significance determined by 1-way ANOVA with Bonferroni posttest comparing bracketed bars; ***p*<0.01.

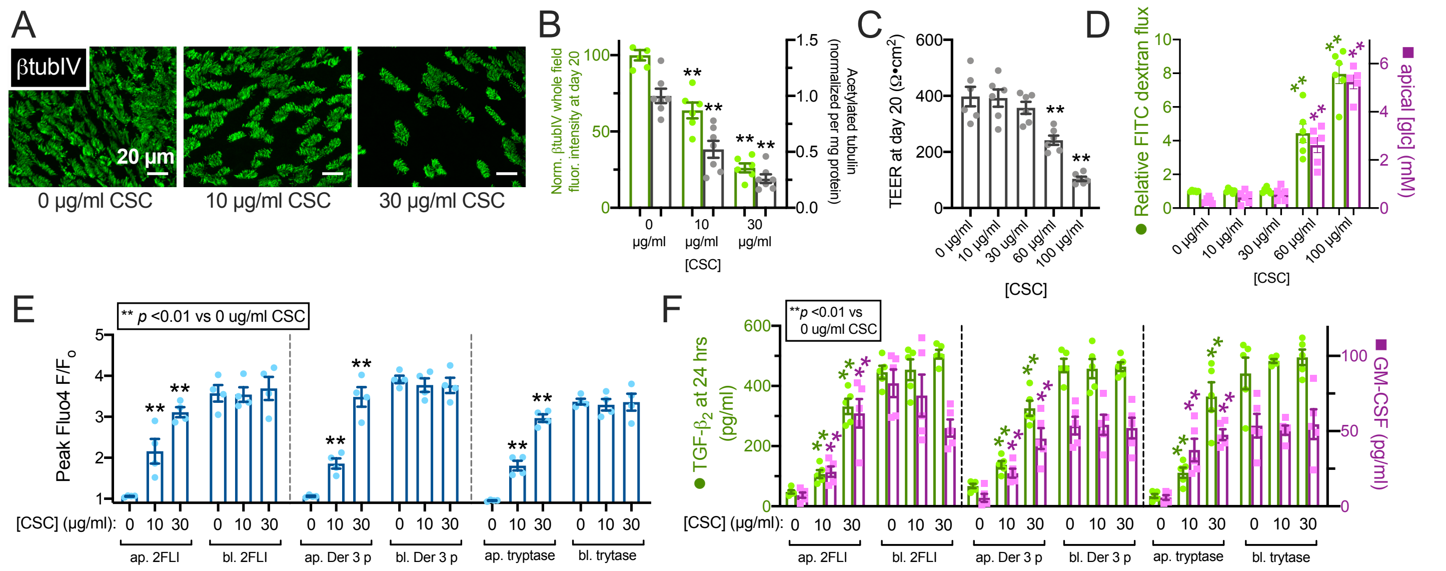


**FIG S11.** Impaired ciliation by cigarette smoke condensate (CSC) exposure alters PAR-2 polarity. **A,** Representative image showing reduction of cilia in ALIs exposed to CSC. **B,** Quantification of β-tubulin IV immunofluorescence (green) and ELISA measurement of acetylated tubulin (gray). **C,** TEER from cultures exposed to CSC for 20 days of ALI differentiation; 30 µg/ml reduced cilia but maintained TEER. **D,** FITC-dextran permeability and apical glucose concentration with CSC, confirming barrier integrity at ≤30 µg/ml CSC. **E,** Peak calcium responses to apical and basolateral 2FLI (25 µM), Der 3 p (1 µM), and tryptase (1 µM) in ALIs with 0, 10, or 30 µg/ml CSC. Note increase of apical 2FLI responses with increased CSC. **F,** TGF-β2 and GM-CSF were measured by ELISA after 24 hours apical or basolateral stimulation as in *E*. Note increased cytokine secretion with apical 2FLI in ALIs exposed to CSC. Significances in *B-G* and *I* determined by 1-way ANOVA with Bonferroni posttest comparing each value to its respective 0 CSC control; ** p<0.01. Data points are from 6-10 individual ALIs from 3-5 human patients.

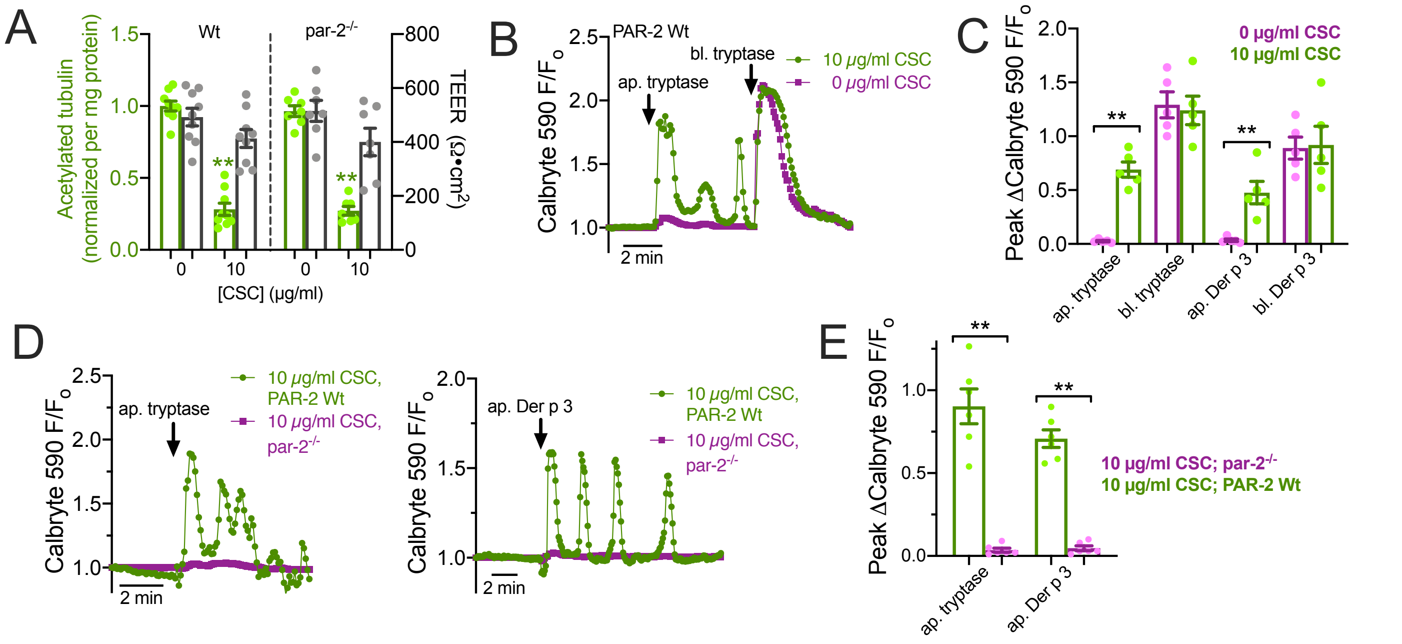


**FIG S12. Verification of PAR-2 polarity alterations by cigarette smoke condensate (CSC) in Wt and par-2$^{-/-}$ mice. A,** Measurement of acetylated tubulin and TEER in mouse nasal ALIs cultured for 20 days ± 10 μg/ml CSC. Acetylated tubulin was significantly decreased with 10 μg/ml CSC but TEER was not. **B,** Representative calcium trace showing apical tryptase (1 μM) response in Wt mouse ALI exposed to 10 μg/ml CSC but not in control ALI (0 μg/ml CSC). **C,** Peak change in Calbryte 590 $F/F_o$ with apical vs basolateral tryptase or Der p 3 (1 μM each) in Wt mouse ALIs exposed to 0 or 10 μg/ml CSC. Note apical responses only observed with 10 μg/ml CSC. **D,** Representative trace of Calbryte 590 response to apical tryptase or Der p 3 (1 μM each) in Wt or PAR-2 knockout (par-2$^{-/-}$) ALIs exposed to 10 μg/ml CSC. **E,** Peak change in Calbryte 590 $F/F_o$ with apical tryptase or Der p 3 in Wt or par-2$^{-/-}$ mouse ALIs exposed to 10 μg/ml CSC. Significance in bar graphs by 1-way ANOVA with Bonferroni posttest comparing Wt vs par-2$^{-/-}$ for each agonist with Data points are from 6-10 individual ALIs from 3-5 mice; **$p$<0.01.

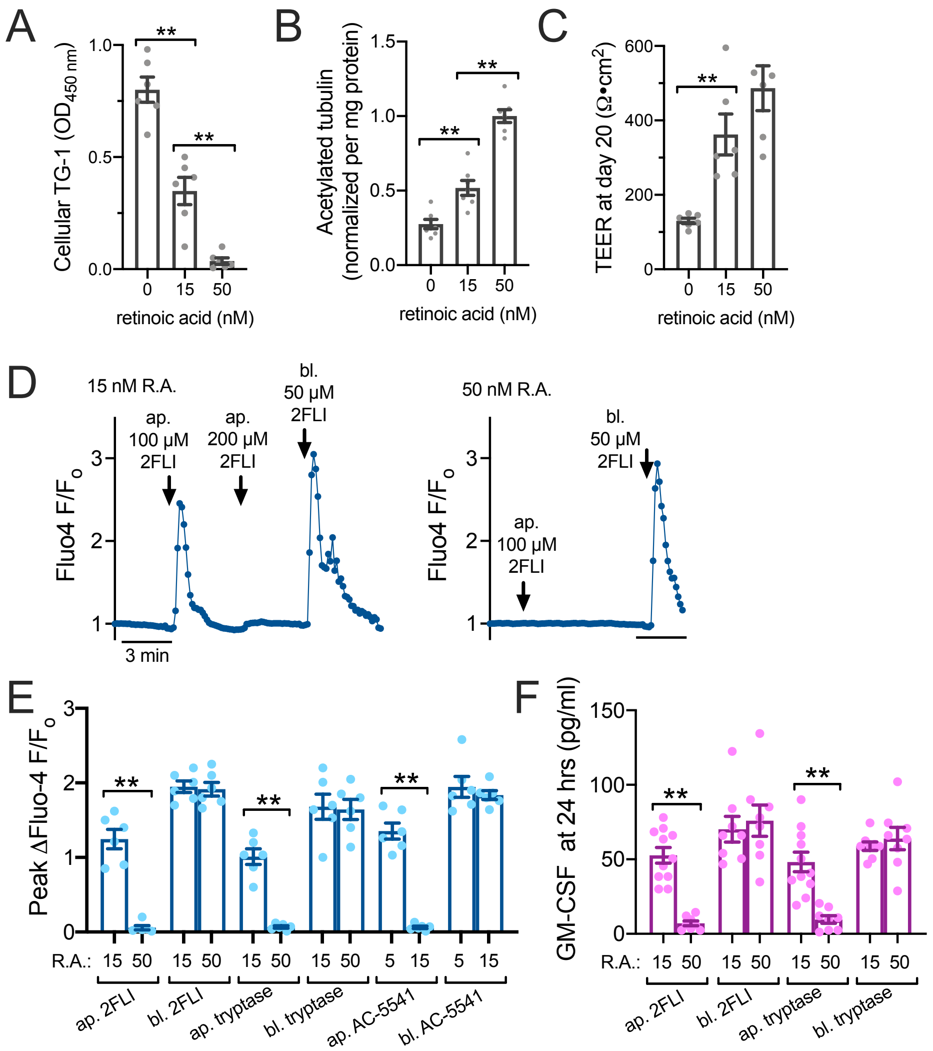


**FIG S13.** Retinoic acid (R.A.) deficiency also alters PAR-2 polarity. **A,** Bar graph showing cellular TG-1 increase in primary human ALIs grown with reduced R.A. **B,** Bar graph showing cellular acetylated tubulin reduction with decreased R.A. **C,** Bar graph showing lower TEER measurements in the absence of R.A. No significant difference was observed between 15 and 50 nM R.A. **D,** Representative Fluo-4 traces showing apical 2FLI response in low R.A. (15 nM) but not normal R.A. (50 nM) culture. Note in left trace that basolateral 2FLI increased calcium even after saturating apical 2FLI, supporting two pools of PAR-2 separated by the epithelial barrier. **E,** Peak change in Fluo-4 F/$F_o$ with apical vs basolateral 2FLI (50 µM), tryptase (10 µM), or PAR-2 agonist AC-5541 (10 µM) in cultures grown in 15 or 50 nM R.A. **F,** GM-CSF was measured by ELISA after apical or basolateral stimulation with 2FLI (10 µM) or tryptase (20 nM). Note responses were observed to apical stimulation in *E* and *F* only in cultures with low R.A. All data points are independent experiments (6-10 per condition) using ALIs cultured from 3-5 patients (2 ALIs per patient). Significance was determined by one-way ANOVA with Bonferroni posttest; $^{**}p<0.01$.
